# Time-averaging and emerging nonergodicity upon resetting of fractional Brownian motion and heterogeneous diffusion processes

**DOI:** 10.1101/2021.04.28.441681

**Authors:** Wei Wang, Andrey G. Cherstvy, Holger Kantz, Ralf Metzler, Igor M. Sokolov

## Abstract

How different are the results of constant-rate resetting of anomalous-diffusion processes in terms of their ensemble-averaged versus time-averaged mean-squared displacements (MSDs versus TAMSDs) and how does the process of stochastic resetting impact nonergodicity? These are the main questions addressed in this study. Specifically, we examine, both analytically and by stochastic simulations, the implications of resetting on the MSD-and TAMSD-based spreading dynamics of fractional Brownian motion (FBM) with a long-time memory, of heterogeneous diffusion processes (HDPs) with a power-law-like space-dependent diffusivity *D*(*x*) = *D*_0_ |*x*| ^*γ*^, and of their “combined” process of HDP-FBM. We find, i.a., that the resetting dynamics of originally ergodic FBM for superdiffusive choices of the Hurst exponent develops distinct disparities in the scaling behavior and magnitudes of the MSDs and mean TAMSDs, indicating so-called weak ergodicity breaking (WEB). For subdiffusive HDPs we also quantify the nonequivalence of the MSD and TAMSD, and additionally observe a new trimodal form of the probability density function (PDF) of particle’ displacements. For all three reset processes (FBM, HDPs, and HDP-FBM) we compute analytically and verify by stochastic computer simulations the short-time (normal and anomalous) MSD and TAMSD asymptotes (making conclusions about WEB) as well as the long-time MSD and TAMSD plateaus, reminiscent of those for “confined” processes. We show that certain characteristics of the reset processes studied are functionally similar, despite the very different stochastic nature of their nonreset variants. Importantly, we discover nonmonotonicity of the ergodicity breaking parameter EB as a function of the resetting rate *r*. For all the reset processes studied, we unveil a pronounced resetting-induced nonergodicity with a maximum of EB at intermediate *r* and EB ∼ (1*/r*)-decay at large *r* values. Together with the emerging MSD-versus-TAMSD disparity, this pronounced *r*-dependence of the EB parameter can be an experimentally testable prediction. We conclude via discussing some implications of our results to experimental systems featuring resetting dynamics.

## I. INTRODUCTION

### A. Overview of recent developments

Resetting a stochastic process, either normal or anomalous^1–10^, via restart events—abrupt or taking a finite time, stochastic/random but distributed, or entirely deterministic in time—returns the particle to its initial position (or a set of positions) according to a certain rule. In recent years, the field of resetting has experienced a wave of new theoretical developments^11–76^ as well as some experimental progress^66,67^. There bouncing-back, often rare, restart events might obey different distributions of waiting times, take place in space-dependent and space-time-coupled manner, with space-time coupled returns, with power-law-like time-dependent resetting rates, with probabilities depending on the offset position, in different geometries and higher dimensions, in comb-like structures, in confining potential wells, under spatial constraints, in nonexponential resetting protocols, with interactions, in coagulation-diffusion processes, in reaction-diffusion protocols, in the presence of space-dependent diffusivity, for the over- and under-damped particle dynamics, to mention a few recent directions of resetting studies.

In most studies, resetting is considered for classical paradigmatic Brownian motion (BM), while resetting studies for more sophisticated fractional- and anomalous-diffusion processes, including continuous-time random walks (CTRWs) and Lévy motion, are less common. Certain first-passage-based, search-related, and search-optimization-like problems involve position restart, with one of the most known manifestations being the minimization of the mean first-passage time (MFPT) to a target at intermediate rates of resetting. A relevant recent study is the first-passage-analysis of a particle in a linear potential with a power-law position-dependent diffusivity, *D*(*x*)∼|*x*| ^61^. Note also that, for resetting rates varying in time, the nonequilibrium stationary state (NESS)^25,77^ was shown^34^ to exist for a decay of *r*(*t*) slower than ∼1*/t*. A stochastic velocity reversion for run-and-tumble particles was also considered^40^.

### B. Some examples of resetting

The list of real-world processes epitomizing resetting/restarting includes, but is not limited to,

- optimization of some foraging or search strategies^78–85^ [employed by animals, fish, insects, microorganisms, bacteria, immune T-cells, etc.] and in behavioral biology^86^ (e.g., by a diffusion process with two distinct modes: with a detailed local search and rapid relocations between “patches” likely to yield food)^87–89^, relocation of animals to previously frequently visited places^20^ and movement-ecology data^90^ in *macrobiological sciences*.
- Some examples from the *nano- and micro-biological* world are the events of stochastic resetting due to backtracking interrupting a processive motion of RNA polymerase along DNA upon transcription^33^ and an alternating switching between the 3D diffusion-based spreading and 1D recognition-based target search in motion of DNA-binding proteins in DNA coils^91–95^.
- On the level of *simple organisms*, stochastic-switching mechanisms between different phenotypes can be mentioned (employed by various bacteria and fungi to optimize adaptation of a phenotypically diverse population of individuals to fluctuating [and, often, irregularly changing] environments^96–101^).
- strategies for boosting combinatorial-search algorithms^102–104^ (via adding a controlled amount of randomization (complete backtracks) to minimize the cumulative search time for a set of tasks or make mean search-times more predictable)^104^ and dependency-directed backtracking algorithms in hard constraint-satisfaction problems involving artificial intelligence^102^ in *computer* sciences,
- in visual pattern recognition, picture-viewing-, and visual-search-strategies^105–109^ (where large-visual-angle jerking-like saccadic motions are interrupted by fixational tremor-like, jiggling microsaccadic “observational” motions [depending, i.a., on the actual task being posed, the contextual information, habituation effects, etc.]^110–115^ and in optimized eye-movement strategies in the brain for visual-search tasks (such as in saccadic models of preferred image-search directions maximizing information about the “target”) in *psychology* ^109,116^.
- In quantitative *financial mathematics*, the models of option pricing for barrier-type and reset-type options (involving option-price adjustments upon crossing certain price boundaries or at preset dates during its life-time) are known for decades^117–121^. Abrupt drops of stock-market prices at times of economic crashes and other sudden catastrophic events^60^ can also be viewed as reset-related phenomena.
- Recent experimental *(wo-)man-made particle-tracking* resetting setups involve a manipulation of micron-sized beads in optical traps^122^ and “tweezers”^66,67^ and can potentially yield time series amenable for a single-trajectory-based time-averaging analysis to decipher the underlying diffusion process, as in the theory developed below.

From the theoretical perspective, the diffusive spread of a stochastic process is in a way “confined” by resetting events yielding in the long-time limit a NESS. In this state, the probability density/distribution function (PDF) is quasi-stationary, but the system still features probability fluxes due to perpetual resetting events of positions of the particles^14^. The main focus of theoretical and simulations-based investigations of resetting in various stochastic processes *so far* was often on the shapes of the PDF (in the NESS and in the particle-displacement phase), scaling relations and plateaus for the mean-squared displacement (MSD), as well as certain first-passage-time-, search-, and surface-adsorption-related quantities.

## II. RESETTING OF SBM: RECENT RESULTS

Recently, the implications of resetting on the behavior of the MSD and PDF of scaled BM (SBM)^7–9,123–131^ with a time-dependent diffusion coefficient of the power-law form,

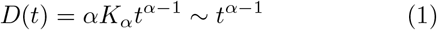

with *α* > 0, was considered both for exponential and power-law distributions of waiting times between two consecutive resetting events. We refer the reader here to the two extensive (mainly MSD- and PDF-focused) studies of Refs.^53,54^. Both nonrenewal (or partial) re-setting setups (with resetting of the position only, while keeping the value and the time-variation of the diffusivity unaltered)^53^ and renewal (complete resetting of particle position and diffusivity)^54^ setups for the SBM diffusivity *D*(*t*) in Eq. (1) were considered.

For nonrenewal exponential resetting of single-particle diffusion, with the waiting-time distribution

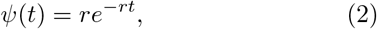

(a Poissonian precess with a constant rate *r* and exponential PDF of waiting times between randomly occurring resetting events), the MSD of reset SBM in the limit of strong resetting and long times, at

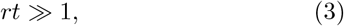

was shown to be^53,54^

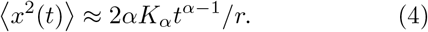

So, the MSD of reset SBM acquires an exponent by one smaller than that of conventional or nonreset SBM, with the MSD

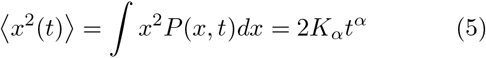

and PDF

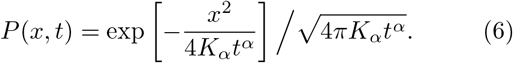

Here *K*_*α*_ is the generalized diffusion coefficient (with the physical units [*K*_*α*_] =m^2^/sec^*α*^) and *α* is the anomalous scaling exponent^3,4,8^. Therefore, resetting leaves superdiffusive SBM with *α* > 2 superdiffusive, while superdiffusive SBM with 1 < *α* < 2 is being converted after resetting into a process with a subdiffusive growth of the MSD, and, lastly, initially subdiffusive SBM with 0 < *α* < 1 gets totally localized by resetting (all particles are accumulated near the origin at long times).

For a power-law-like (non-Poissonian) nonrenewal resetting, with *ψ*(*t*) = (*β/τ*_0_)*/*(1 + *t/τ*_0_)^1+*β*^ and *β* > 0, it was shown^53,54^ that the MSD for reset SBM keeps the exponent of basal SBM and gets only reduced in magnitude for 0 < *β* < 1. In the range *β* > 2 the scaling exponent of the MSD of SBM with resetting gets reduced by one, similarly to the case of exponential or Poissonian resetting in Eq. (4), so that^53,54^ ⟨*x*^2^(*t*) ⟩ = 2*ατ*_0_*K*_*α*_*t*^*α*−1^*/*(*β* ^−^ 2).

For renewal resetting (when both the particle position and diffusivity are being reset), for SBM with exponential resetting the MSD was shown to approach a NESS plateau with the saturation level at^53,54^

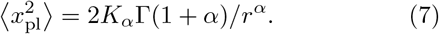

This turns into the known result for BM with exponential constant-rate resetting, namely

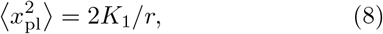

where Γ(*y*) is the standard Euler-Legendre Gamma function.

For a power-law resetting with 0 < *β* < 1, in contrast, the MSD of reset SBM keeps the scaling exponent of free SBM, while the MSD magnitude gets altered by a factor containing the waiting-time PDF exponent *β*, namely^53,54^ ⟨*x*^2^(*t*)⟩ = 2*K*_*α*_*t*^*α*^ × Γ(*α*−*β*+1)*/*[Γ(*α*+1)Γ(1−*β*)]. The MSD behavior differs dramatically for scaling exponents in the region 1 < *β* < 2, namely at long times for *β* < *α* + 1 the MSD scales as ⟨*x*^2^(*t*) ⟩ ∼ *t*^*α*+1−*β*^, while for *β* > *α* + 1 the MSD approaches a long-time plateau with *α*- and *β*-dependent values.

For CTRWs, a similar type of the MSD- and PDF-based analysis for complete resetting (of particle positions and of waiting times) and in the case of partial resetting (of particle positions only) was executed as well^55^. For complete resetting, for CTRWs the behaviors of the MSD and PDF were shown to be the same as for the corresponding SBM. This is expected because SBM is known to be the mean-field model of CTRWs^124^.

## III. OUTLINE OF THE PAPER

Our main objective here is to enrich the list of important physical observables employed in resetting-dynamics studies by a single-trajectory-based time-averaged MSD (TAMSD) and the ergodicity breaking parameter EB, see Eqs. (9) and (11) below. The TAMSD is a quantifier often implemented in single-particle-tracking experiments and its characteristic features have been intensely developed theoretically over the last years for a variety of non-reset stochastic processes featuring anomalous diffusion^8^. The EB parameter characterizes the spread of individual TAMSDs and describes the degree of nonergodicity^8^.

The primary focus of the current study is on the effects of resetting onto the TAMSD characteristics for stochastic processes of nearly ergodic (see Refs.^132–135^) fractional BM (FBM)^42,136–144^and of nonergodic heterogeneous diffusion processes (HDPs)^126,143,146–150^ as well as for a combination of FBM with varying Hurst exponent *H*^151,152^ and HDPs with varying exponent of the power-law-like diffusivity *γ*^150^. Note that a “hybrid” process of SBM-HDP was also introduced^126^ and recently applied to the experimental data^153^. The implications of resetting onto the MSD and PDF of FBM, HDPs, and HDP-FBM are also considered, being a secondary focus.

Note that, despite identical PDFs of particle displacements governing SBM and FBM, SBM is a memoryless Markovian process featuring nonstationary increments and nonequivalence of the MSD and TAMSD^128^ (often also indicative of weak ergodicity breaking, WEB), whereas, in contrast to SBM, FBM is an innately non-Markovian process with a long-time memory and stationary displacement increments, for which the MSD and mean TAMSD are [statistically] equivalent (considered as an ergodic process in this sense^8,139,144^). The standard quantifier of ergodicity—the so-called ergodicity breaking parameter denoted as EB in Eq. (11) below—behaves for SBM and FBM, however, rather similar in terms of EB approach to zero at vanishing lag times and for long trajectories, at Δ*/T* ≪ 1. Generally, the stationarity of increments of a stochastic process is a prerequisite of its ergodicity.

The paper is structured as follows. We define the observables and present the details of the simulation scheme in Sec. IV. In Sec. V the results for the MSD and mean TAMSD for the resetting dynamics of FBM are presented, as obtained from stochastic computer simulations. In Secs. VI and VII we examine the implications of resetting onto the MSD, PDF, and mean TAMSD of HDPs and HDP-FBM, respectively. The results for the resetting dynamics of FBM and HDPs are compared to those for SBM (outlined in Sec. II). We thus start from the known results for reset SBM in Sec. II, move to reset FBM (examined now in terms of the MSD, PDF, TAMSD, and EB) featuring some commonalities with reset SBM, and, lastly, enter the complete *terra incognita* of reset HDPs and HDP-FBM processes, again, examined these reset stochastic processes with the standard measures (MSD and PDF) and novel quantifiers (TAMSD and EB). Finally, we draw conclusions and discuss some applications of our results in Sec. VIII.

## IV. OBSERVABLES, MODELS, AND SIMULATION SCHEME

### A. Definitions of physical observables

We employ the concept of single-trajectory-based averaging along the time series of particle positions *x*(*t*) in terms of the TAMSD^8^

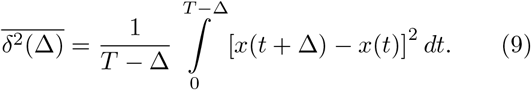

Here, Δ is the so-called lag time and *T* is the total length of the time series. After averaging over *N* statistically independent TAMSD realizations, the mean TAMSD is computed as the arithmetic mean

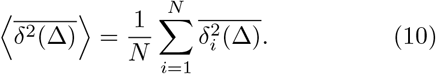

The angular brackets denote hereafter averaging over realizations of noise, while averaging over time is denoted by the overline.

To quantify the degree of ergodicity, the so-called ergodicity breaking parameter EB is utilized^8,139,145^

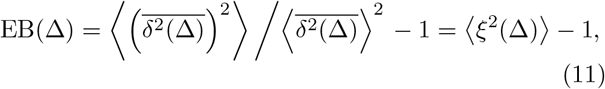

where the dimensionless variable

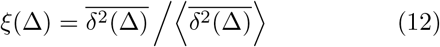

describes the dispersion of individual TAMSD realizations around their mean (10). The distribution of TAMSDs for an ensemble of particle trajectories is characterized by the PDF *ϕ*(*ξ*)^8,143^.

### B. Diffusion models

#### 1. Main equations for FBM, HDPs, and HDP-FBM

We employ the same simulation scheme as in the recent study of a numerical and analytical investigation of the “compound” process of FBM-HDPs introduced recently in Ref.^150^. Shortly, the overdamped Langevin equation

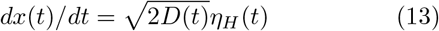

with fractional Gaussian zero-mean noise *η*_*H*_ (*t*) featuring the power-law correlation function (for *t* ≠*t* ^′^),

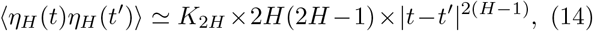

is used to simulate free^136,137^ and reset FBM in terms of single-particle diffusion.

For HDPs, the same Langevin equation for a zero-mass particle,

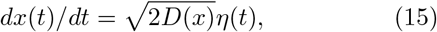

is modeled with white Gaussian noise having zero mean and unit variance,

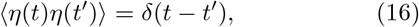

and position-dependent diffusion coefficient of a power-law form^146^,

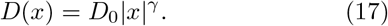

For *γ* < 0, in order to regularize the diverging diffusivity at the origin, *x* = 0, a modified position-dependent diffusion coefficient is used in simulations, namely^146,150^

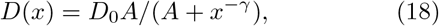

with the factor *A* = 10^−2^ [acting as a small offset]. For 2 > *γ* > 0 no problems with diverging diffusivities occurs and the form (17) is used directly. The critical point for the diffusivity exponent is at *γ* = 2: upon approaching this point the exponential (and not a power-law-like) growth of the MSD is realized^147,154^.

For a stochastic process of HDP-FBM we consider analytically and simulate the overdamped Langevin equation

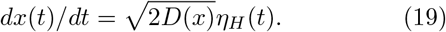

Not to be repetitive, we refer the reader to Ref.^150^for the the details of the analytical solutions of these equations, the description of the region of parameters of FBM and HDPs amenable for a solution, etc.

#### 2. Some applications of FBM and HDPs

The model of FBM was recently applied to rationalize (•) the non-Brownian anomalous dynamics of lipids, cholesterols, and proteins in/on lipid membranes^155–158^, (•) the dynamics of G-proteins and G-protein-coupled receptors on plasma membrane^159,160^, (•) the diffusion of labeled mRNAs in living bacterial cells^161^, (•) anomalous motion of lipid granules in living yeast cells^162^, (•) the diffusion of telomeres inside the nuclei of human cancer cells^163,164^, (•) the dynamics of chromosomal loci in bacterial cells^165–167^, (•) non-Gaussian nonergodic anomalous diffusion of micron-sized beads in mucin-polymer hydrogels^168^, (•) tracer dynamics in actin networks^169^ [for the particles larger in size than the network meshsize]^170^, (•) heterogeneous intracellular transport of DNA cargo in cancerous cells [with coexisting ergodic-and-nonergodic but nonaging dynamics]^171^, as well as (•) intermittent bulk-surface non-Gaussian and aging anomalous diffusion with aging of anticancer-drug doxorubicin in silica nanoslits^172^, to mention a few examples.

Solutions of the diffusion equation with variable diffusion coefficients go back to Boltzmann^173^, while the nonlinear diffusion equation with the diffusivity being a power law of concentration of the diffusing substance,

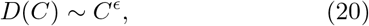

with *ϵ* > 0, was solved by Pattle^174^ (see also some recent “reincarnations”^175,176^). Contemporary models of diffusion with space-dependent diffusion coefficients^154,177–184^—with HDPs being a specific example that assumes the functional diffusivity form (17)— can be used to describe (•) the non-Brownian diffusion in crowded, porous, and heterogeneous media^185–202^ (such as densely macromolecularly crowded cell cytoplasm), (•) the reduction of a critical “patch size” required for survival of a population in the case of heterogeneous diffusion of its individuals^181^, (•) diffusion in heterogeneous comb-like and fractal structures^182^, (•) escalated polymerization of RNA nucleotides by a spatially confined thermal (and diffusivity) gradient in thermophoresis setups^203^, (•) motion of active particles with space-dependent friction in potentials [both of power-law forms]^204^, and (•) transient subdiffusion in disordered space-inhomogeneous quantum walks^205,206^. We mention also a class of diffusion models with (•) particle-spreading scenarios with concentration-dependent power-law-like diffusivity (20)^175,207^, (•) concentration-dependent dispersion in the population dynamics, with a nonlinear dependence of mobility on particle density, *D*(*ρ*) ∼ *ρ*^*κ*^ (yielding a migration from more- to less-populated areas)^208–210^, as well as (•) similar nonlinear equations^10^ for porous-media dynamics^211^, nonlinear heat-conductance systems (with a power-law-like temperature-dependent conductivity), and the dynamics of granular materials^211,212^.

### C. Implementation of resetting: algorithm of numerical simulation

We use below the exact theoretical results and asymptotic relations of Ref.^150^ for FBM, HDPs, and HDP-FBM processes (without re-deriving them here) and employ the middle-point, physically motivated, Stratonovich-convention-based simulation scheme for Eqs. (13), (15), and (19) (see the details of the Itô-Stratonovich conversion employed and Eq. (32) in Ref.^150^). For all these processes at each elementary time-step of *dt* = 10^−2^ or *dt* = 10^−3^ in simulations a constant rate of resetting is set *r*, so that resetting probability to the initial position

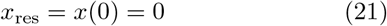

is *r* × *dt*, see Fig. 1. When simulating reset FBM we always employ (21), while for reset HDPs and HDP-FBM an offset *x*_res_ = 0.01 is used (to avoid stalling of particles at the origin, in particular for superdiffusive HDPs).

**FIG. 1:**
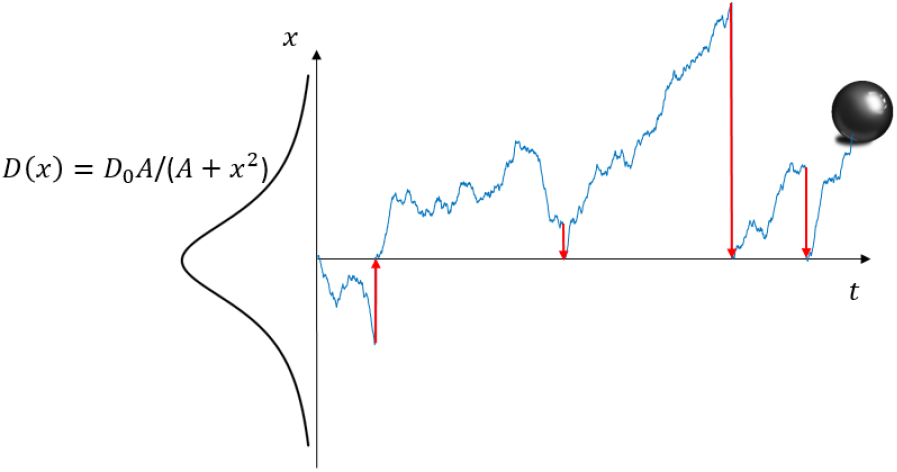
Simulated trajectory of a subdiffusive HDP (18), with the diffusion-coefficient *D*(*x*) ∼|*x*| ^−2^ being shown, in the presence of Poissonian resetting of particles to *x* = 0.

The waiting-time distribution of resetting events (2) yields the average resetting time

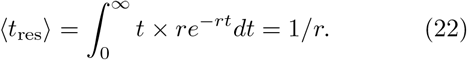

We consider instantaneous resetting or “jumping-to-the-origin” events that makes a consistent picture with the inertia-free or overdamped dynamics of the particles in their displacement phase (between the events of resetting). Other-than-Poissonian distributions of resetting times, noninstantaneous resetting protocols (constant-velocity, etc.), resetting triggered by crossing of certain thresholds, etc. can also be considered and implemented in simulations.

Resetting of FBM, similarly to that of SBM^53,54^, can be performed in a partially or fully renewal scheme. In the second case, the memory of noise correlations (14) is completely “erased” upon each resetting event: we use this *fully renewal scheme* when simulating exponentially reset FBM and HDP-FBM below. For HDPs, the implementation of resetting is unproblematic because the process is memoryless. In simulations, we generate particle trajectories for all three reset processes considered via a discretization scheme with a Poissonian-resetting protocol: the particle position *x*(*t* +*dt*) is generated depending on the previous point *x*(*t*) via the standard Langevin-dynamics simulation approaches (using Eqs. (13), (15), and (19)) when no resetting occur during the time *t* + *dt* [with the probability 1 −*r* ×*dt*] and the particle is moved to the reset position *x*_res_ [with the probability *r* × *dt*] when the resetting event does take place.

## V. RESETTING OF FBM

For reset FBM with a fully erased memory—as well as for other Gaussian (Markovian as well as non-Markovian) stochastic processes with the same free-motion PDF as that of SBM—it was predicted (provided after each resetting the particle performs statistically identical diffusion process) to have the same MSD and PDF as for renewal resetting of SBM^54^. In general, one can expect that, both for FBM and HDPs, the events of instantaneous resetting of particle positions to the origin will give rise to *larger* displacements for consecutive time-steps. This not only contributes to a growing magnitude of the TAMSD at short lag times (see below), but also can give rise to a more pronounced irreproducibility of individual TAMSD trajectories and, therefore, larger values of the EB parameter in this region. In the limit of long time, in the NESS the MSD and TAMSD are expected to stagnate and lose any dependence in their growth with (lag) time. We substantiate on these intuitive expectations below.

### A. MSD

We present the results of analytical computations and findings of computer simulations for the MSD of reset FBM in Fig. 2 (also showing the TAMSD realizations), for several values of the Hurst exponent *H*, for both sub- and superdiffusive dynamics (in terms of the MSD). For smaller values of *H* we used a smaller time-step of *dt* = 10^−3^ in order to better resolve the short-time behavior of the ensemble- and time-averages. The MSD of FBM with exponential resetting starts unperturbed with the expected short-time behavior characteristic of anomalous diffusion^8,139–141^, namely

**FIG. 2:**
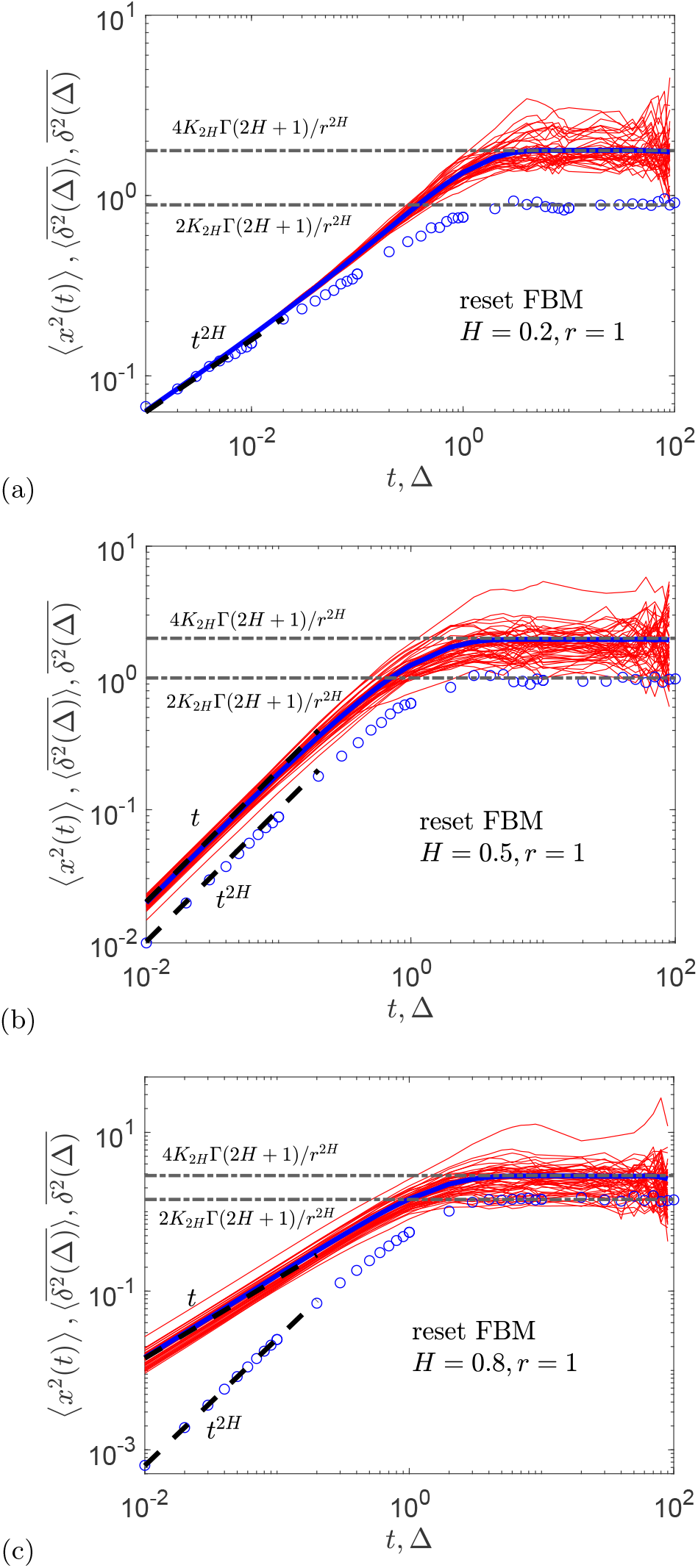
Magnitude of the MSD (blue circles), the spread of individual TAMSDs (thin red curves), and the mean TAMSD (thick blue curve) for the dynamics of reset FBM, shown for varying values of the Hurst exponent (the values of *H* are indicated in the graphs). Simulations of resetting are conducted with no FBM-related memory effects. Theoretical long-time plateaus for the MSD and mean TAMSD are given by Eqs. (25) and (32), respectively. The short-time asymptote for the MSD is Eq. (23), while the evolution of the mean TAMSD at short lag times follows Eqs. (30) and (32), for subdiffusive and superdiffusive FBM, correspondingly. The asymptotes are shown as the black dashed and dot-dashed lines. Parameters: the length of the trajectories is *T* = 10^2^, the elementary time-step in simulations is *dt* = 10^−2^ (except for *H* = 0.2 when *dt* = 10^−3^), the number of trajectories for ensemble averaging is *N* = 10^4^, the resetting rate is *r* = 1 [the resetting probability per step is *r* × *dt* = 10^−2^], and the generalized diffusion coefficient is set to *K*_2*H*_ = 1*/*2.

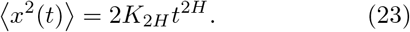

At times of the order of the average resetting time (22),

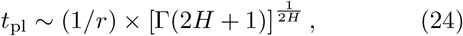

the MSD starts saturating at a plateau (with plateau-related quantities denoted by the subscript “pl” hereafter), with the same height as that for reset SBM^53,54^, see Eq, (7). Specifically, for NESS the height of the MSD plateau (with a substitution *α* = 2*H*) is

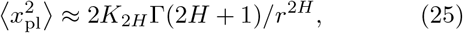

as indicated in Fig. 2. We have checked that this transition behavior for the MSD is the same for the scenarios with and without long-time memory of FBM (14) being included in simulations (results not shown). The consistency of the short-time MSD asymptote (23), the *r*-dependent MSD plateaus (25), and different onset times onto the NESS-related MSD behavior is demonstrated in Figs. S1 and S2 via presenting the results of computer simulations and the theoretical predictions for a set of several rates of resetting *r*.

### B. PDF

The results for the PDF at intermediate-to-large displacements for reset FBM in the NESS are in full agreement with the predictions for fully-reset SBM, given by a time-independent function of the form (with the substitution *α* = 2*H* to go from SBM- to FBM-expressions)^53,54^

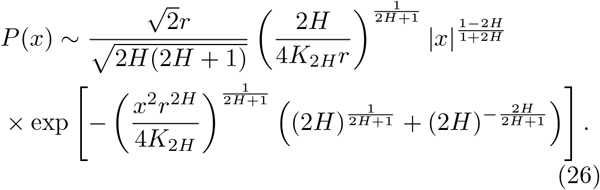

The leading functional behavior for the PDF “tails” in expression (26) is a stretched-or-compressed exponential function, namely^53,54^

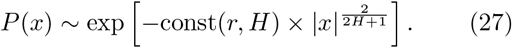

We have checked that the levels for the MSD plateaus in the NESS, 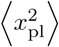, are well described using the approximate PDF (26), especially at Hurst exponents *H* ≳1/2, as shown in Fig. S4.

In Fig. 3 we present the agreement of the results of computer simulations for the PDF of reset FBM with the theoretically shape (26) at long times in the NESS and for intermediate-to-large displacements of the particles. The shapes of the PDF for reset FBM with and without memory are statistically identical at these conditions. We prove this in Fig. 3 presenting the results for reset FBMs for sub- and superdiffusive choices of *H*. Note also that the MSD evolution for reset FBM with and without memory is also identical (results not shown). For resetting rate *r* = 1 the FBM dynamics at *H* = 0.8 is much faster than at *H* = 0.2. For a preset/fixed length of simulated trajectories, the MSD plateaus for reset FBM at *H* = 0.8 occupy a rather extended time domain, while the quasi-stationary state for a slower dynamics at *H* = 0.2 is not yet realized, compare Figs. 2c and S2b at *r* = 1.

**FIG. 3:**
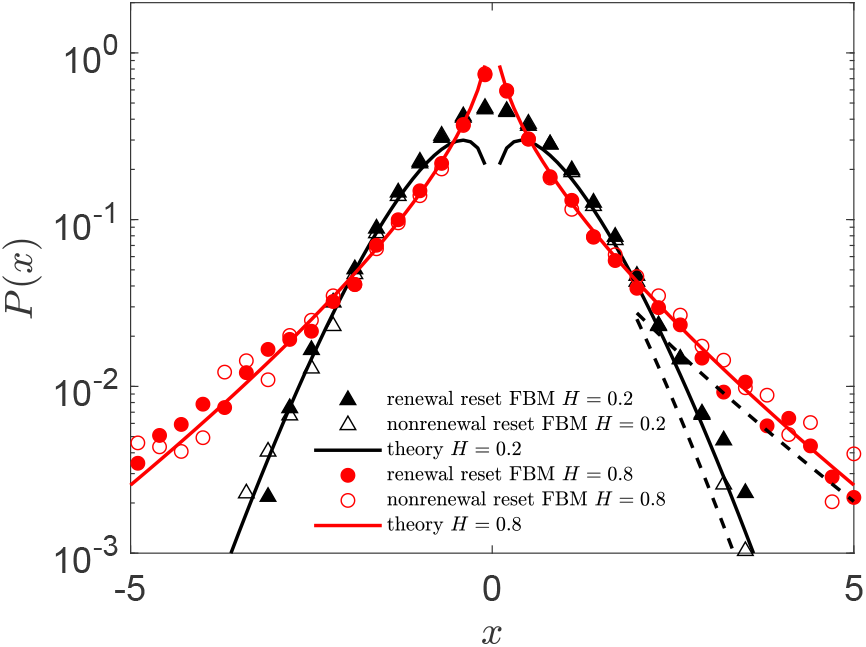
Shapes of the PDFs of reset FBM, computed for the Hurst exponents *H* = 0.2 with *dt* = 10^−3^ and *H* = 0.8 with *dt* = 10^−2^ (see the legend) at *r* = 1. The symbols represent the results of simulations, while the solid curves are the theoretical intermediate-to-long-times expectations for the PDF given by (26). The dashed lines at large displacements are stretched- or compressed-exponential asymptotes for the PDF tails^53,54^ given by (27).

The PDF shapes for reset FBM evolve upon approaching the NESS. We find that for *r* = 1 the PDF shape at the origin for strongly subdiffusive reset FBM is almost smooth and Gaussian (as for nonreset FBM), while for the same rate of resetting for strongly superdiffusive reset FBM the PDF exhibits a pronounced cusp at the origin stemming from the returning events of the particles. This cusp is well described by the theoretical prediction (26), see Fig. 3, and it corroborates with the emergence of the NESS plateaus of the MSD and mean TAMSD. The empty and filled symbols in Fig. 3 show the results of reset-FBM simulations with and without the long-time memory of noise being taken into account: we check the validity of the theoretical PDF in Fig. 3 once for nonrenewal resetting of FBM too. Therefore, for considerably larger values of *r*, when for strongly subdiffusive reset FBM the MSD also reveals a plateau at long times in the NESS (results not shown, but see Fig. S2c for the onset on this saturating behavior at *r* = 3), the respective PDF of reset FBM at *H* = 0.2 also has a cusp at *x* = 0, as shown in Fig. S5.

The PDF cusp for subdiffusive reset FBM is not in agreement with (26) and it critically depends on the simulation time-step. Ideally, the probability of resetting each step should be small in order for the results to be independent on the time-step used in simulations. The numerical integration of the long-time limit of the PDF (B1)—evidently step-size-independent—does not reveal any cusp at *x* = 0, see the cross-symbols in Figs. S5 and S6. As we reduce the simulation time-step from *dt* = 0.01 to 0.001 keeping the resetting rate the same, the PDF peak in simulations for *H* = 0.2 vanishes and the PDF form nicely agrees with the results of numerical integration of (B1), see Fig. S6. We stress therefore that a sufficient care and accuracy should be taken when simulating such processes.

This numerical inaccuracy giving rise to the “spurious” PDF cusp for *H* = 0.2 gives rise to small, but systematically measurable, deviations in the plateau heights of the MSD and mean TAMSD in the NESS for the simulations for very large resetting rates *r* (results not shown). Note that for reset SBM^53,54^ the PDF shape from the theory was shown as a combination of the results of the Laplace approximation (26) [valid at intermediate-to-large separations from the origin], while the central PDF part near *x* = 0 was the cusp-free numerical integration of the exact PDF expression in Eq. (B1).

Note that the approximate Laplace-method-based PDF (26) is generally not normalized: for subdiffusive (superdiffusive) rest FBM it underestimates (overestimates) the integral ∫ *P* (*x*)*dx*, as shown in Fig. S7. We still call this approximate non-normalized distribution-function a PDF hereafter, for brevity. The PDF (26) describes the simulation data well at intermediate-to-large separations^53,54^, for |*x*| ≳ *x*^*\**^. The deviations from the results of computer simulation at small |*x*| values from the expression (26) are especially pronounced for very subdiffusive reset FBM. A rough estimation for the threshold separation *x*^*\**^ for reset FBM with *H* < 1*/*2 follows [based on the realized PDF shapes] from solving 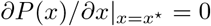 for the inflection point, that gives

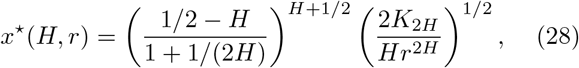

see the inset of Fig. S7 showing *x*^*\**^(*H*) variations. The exponential decay of the PDF of reset FBM in the NESS given by (27) [identical to that of full expression (26)] as well as the scaling relation for the growth-dynamics of the NESS domain with time given by ∼ *t*^*H*+1*/*2^ were obtained before in Ref.^25^. The NESS starts getting established from the restart position [by multiply reset trajectories/particles], with the spatial NESS domain growing quicker in time than a typical FBM diffusion length that scales as 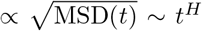. Outside of this PDF- and MSD-stationarity domain, the reset system still performs relaxation and features a time-dependent PDF [describing the trajectories with almost no resetting occurred so far]. In Fig. S8 the PDF decay derived in Ref.^25^ is explicitly compared to our simulation data for subdiffusive reset FBM.

To quantify the height of the PDF at the point of particle resetting in the NESS, in Fig. S9 we show the results of computer simulations for the values of *P* (*x* = 0) for reset FBM. As for nonreset FBM, the PDF is identical to that of pure SBM (given by Eq. (6)), using the PDF-transformation relation (B1), for reset FBM in the NESS the PDF at the point of particles’ return assumes the value

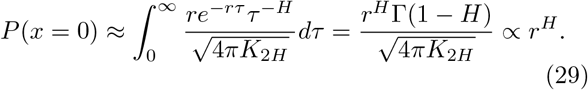

The asymptotic long-time law *P* (*x* = 0) ∼ *r*^*H*^ excellently agrees with the results of our simulations, see Fig. S9.

### C. TAMSD

We find from simulations that for subdiffusive Hurst exponents, with 0 < *H* < 1*/*2, the TAMSD starts sublinearly and has roughly the same magnitude as the short-time MSD (23), namely

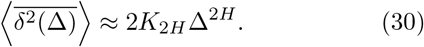

For initially superdiffusive FBMs, in the range of Hurst exponents 1*/*2 < *H* < 1, in contrast, the mean TAMSD is linear at short lag times,

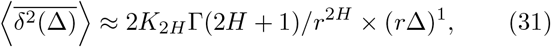

as shown in Fig. 2. In App. A we provide the derivation of asymptotes (30) and (31). For *H* = 1*/*2 we need to take the sum of two independent terms (30) and (31) in order to *quantitatively* fit the short-lag-time behavior of the mean TAMSD.

At long lag times for reset FBM the TAMSD plateau in the NESS is realized, with the magnitude of twice that of the MSD plateau (given by Eq. (25)), namely

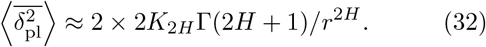

The ratio

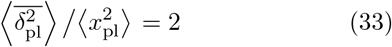

in the long-time (quasi-stationary) limit is known also for FBM confined in harmonic potentials^141^ and interval-confined HDPs^148^. This twice-the-MSD magnitude in (32) stems from the TAMSD definition (9) and is not related to resetting *per se*. The reason being that after multiple resettings in the NESS the process values at *x*(*t* + Δ) and *x*(*t*) become almost independent, so that

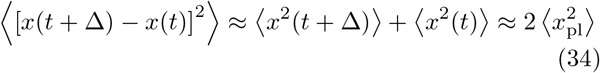

yielding (33).

We stress, however, that for FBM in parabolic potentials at short lag times the MSD and mean TAMSD are fully equivalent in magnitude and scaling^141^. This fact is in stark contrast to reset FBM studied here, where WEB and MSD-versus-TAMSD nonequivalence emerges at *H* > 1*/*2, see also Tab. I. Indeed, we observe that for reset FBM the mean TAMSD in the region of short lag times always increases in magnitude as compared to that of free/nonreset FBM. At the very last point of the trajectory, at Δ → *T*, the magnitude of the mean TAMSD approaches that of the MSD in the plateau region. This fact is, however, not very well visible in the presentation in Figs. 2, S1, and S2 because of a logarithmic sampling of the simulation data (with only ten points per decade).

Via equating the magnitudes of the short-lag-time TAMSD asymptotes and the long-time TAMSD plateau one can assess the lag time at which the TAMSD plateau (32) starts to be followed as 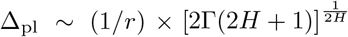 for 0 < *H* < 1*/*2 and Δ_pl_ ∼ 2*/r* for 1*/*2 < *H* < 1.

To assess the effects of a varying resetting rate *r*, in Fig. S1 we present the results of computer simulations for the largest Hurst exponent *H* = 0.99 (when the scatter of individual 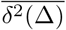 trajectories is the broadest [for the same (fixed) *r*], see also Sec. V D) at varying *r* values. We observe that the predicted *r*-dependent plateaus for the MSD and mean TAMSD at long times in the NESS, expressions (25) and (32), respectively, excellently describe the results of simulations.

The spread of individual TAMSD realizations for the resetting dynamics of FBM typically increases, as compared to that of free FBM that a process with EB(Δ) → 0 for long trajectories and short lag times (at Δ*/T* → 0) in the continuous-time formulation (see Refs.^132–134^ for the definition(s) and general discussion of ergodicity), as demonstrated in Refs.^8,139,143,144^. This effect is particularly pronounced for large superdiffusive Hurst exponents, *H*, as illustrated in Figs. 2c and S1. In the region of resetting rates *r* considered in Fig. S1 for *H* = 0.99 the relative spread of the TAMSDs increases with decreasing *r*. Note, however, that the entire variation of the dispersion of individual TAMSDs as a function of the Hurst exponent and resetting rate is rather nontrivial, as we unveil below.

In contrast, for strongly subdiffusive Hurst exponents, see, e.g., the results presented in Fig. S2 for the choice *H* = 0.2, the impact of the resetting rate [varying in the same interval] is rather weak. This is intuitively clear: as compared to the trajectories of strongly superdiffusive FBM which depart far away from the origin and thus are strongly impact by a given resetting rate, for very subdiffusive FBM the trajectories are weakly fluctuating, meandering in a close proximity of the starting position, so that the impact of events of particle’s resetting to zero is nearly unnoticed in our quantifiers. To substantiate on this claim, in Fig. S3 two FBM trajectories for each choice of *H* = 0.8 and *H* =0.2 are presented in the absence and in the presence of resetting.

### D. EB

#### 1. Distribution of TAMSDs

The distribution of TAMSDs for *H* = 0.8 for FBM in the presence of resetting is presented in Fig. S10. We find that for small rate of resetting the distribution *ϕ*(*ξ*) at short lag times for a weakly reset FBM is considerably skewed toward the region *ξ* > 1. This fact (known not only for FBM^143,146^) stems from the existence of a natural boundary at *ξ* = 0 (as *ξ* is positively defined, Eq. (12)) and an unbounded domain extending to *ξ* ≫ 1. As the rate of resetting increases (in the range of rates chosen), the TAMSD realizations become progressively less scattered around their mean, indicative of a more reproducible [or ergodic] dynamics in terms of scatter of individual TAMSD magnitudes, described by *ϕ*(*ξ*).

#### 2. Large r values

This enhanced reproducibility of TAMSD realizations in this range of *r* is also reflected in the decreasing value of the EB parameter, see the simulation data in Fig. 4 for *H* = 0.8. A qualitatively similar behavior of EB versus *r* is also observed in simulations for *H* = 1*/*2, but in a smaller range of resetting rates, see Fig. 4. At these “intermediate” *r* values for reset FBM with elevated Hurst exponents, the value of the EB parameter at short lag times drops according to

**FIG. 4:**
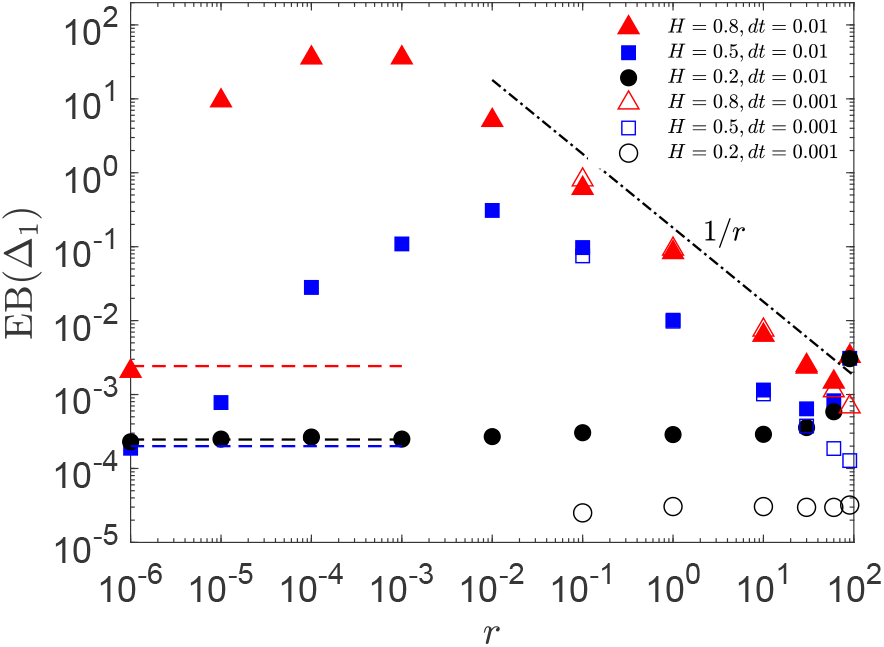
Dependence of the EB parameter (11) on the rate of FBM resetting *r*, computed at Δ = Δ_1_ = 10^−2^ and *dt* = 10^−2^ (filled symbols) and *dt* = 10^−3^ (empty symbols), for a set of *H* exponents. The theoretical asymptote (35) is the black dot-dashed line at intermediate-to-large resetting rates. The values of EB for nonreset FBM for the time-step *dt* = 10^−2^ are the dashed plateaus (with the *H*-respective colors, see also the legend). The length of the trajectories is *T* = 10^2^.

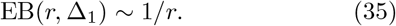

The value of EB itself is also sensitive to *H* values: larger Hurst exponents yield larger EB, as our simulations show.

Therefore, the dependence (35) of EB on *r* in this range of resetting rates for reset FBM with elevated *H* is functionally similar to that of EB on 1*/T* (considered, again, at short lag times) repeatedly detected for a number of (both normal and anomalous) stochastic processes^8^. The latter indicates progressively more ergodic behavior for longer trajectories, so that the relation EB(*T*, Δ_1_) ∼ 1*/T* holds. The decrease of EB values with *r* (in the limit of frequent resetting) for reset FBM with larger Hurst exponents is corroborated by a shrinking *ϕ*(*ξ*(Δ_1_)) distribution computed for the same conditions with increasing *r*, as shown in Fig. S10. In contrast, for strongly subdiffusive reset FBMs we observe a nearly *constant* EB upon varying *r*, as illustrated in Fig. 4 for *H* = 0.2.

For strong or frequent resetting, as the probability of a reset even at each displacement step approaches unity, the EB parameter reveals a sensitivity to the step-size value (discreteness effects), alike those for the emergence of PDF cusps at *x* → 0 in the same limit. For instance, our simulations for a fixed trace length *T* = 10^2^ performed for *dt* = 10^−2^ yield somewhat different large-*r* behavior of EB for superdiffusive reset FBM, as compared to that at *dt* = 10^−3^ (effectively, 10-times more points in each trajectory). Namely, for smaller time-steps the deviations from the expected EB versus *r* scaling (35) disappear, see the empty symbols at large *r* in Fig. 4.

We emphasize here also the essential differences in the behavior and magnitudes of the EB parameter obtained from the discrete-time stochastic simulations versus those predicted from a continuous-time theory of (non-)ergodicity of FBM^139^, as studied in Ref.^144^. For instance, for subdiffusive FBM the magnitude of EB at a fixed lag time Δ_1_ and fixed trajectory length *T* loses its dependence on the Hurst exponent and EB stagnates at a plateau, with the height dropping as

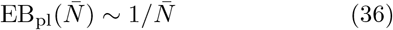

with the number of points in the trajectory, 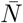. In such a case, the EB parameter scales with the time-step 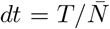 used in simulations. In Fig. 4 the simulation data for reset FBM at *H* = 0.2 for two different time-steps illustrate the EB(*dt*) variation (36), see the empty symbols at large *r* values.

#### 3. Small r values

For very weak or rare resetting, we recover the small values of EB characteristic for ergodic free FBM^8,139,144^ at Δ*/T* ≪ 1 at the corresponding *H* values, computed recently in Ref.^150^ and denoted as the dashed lines of respective colors in Fig. 4. We emphasize, however, that the approach of EB of reset FBM to that of non-reset FBM yields a *nonmonotonic dependence* of EB(*r*), particularly pronounced for strongly superdiffusive (but also present for slightly subdiffusive) reset FBM. This nonmonotonicity of EB(Δ_1_) yields a strongly resetting-enhanced dispersion of short-lag-time magnitudes of individual TAMSDs of reset FBM at intermediate resetting rates. The maximum of EB as a function of *r*— characterizing the strongest irreproducibility of individual TAMSDs—shifts towards smaller *r* values for more superdiffusive FBMs, compare the data for EB(*r*) for different *H* in Fig. 4. We find that the maximally achievable EB values for strongly superdiffusive reset FBM at intermediate *r* are colossal, about four orders of magnitude larger than the respective EB values for free FBM^139,144^.

We also emphasize that for reset FBM the variation of EB(Δ_1_) with resetting rate *r* stays qualitatively similar also for longer trajectories, with EB values getting reduced according to EB(Δ_1_, *T*) ∼ 1*/T* relation (similar to that for free BM^8^ and free FBM with 0 < *H* < 3*/*4^139,144^), see Fig. S11.

## IV. RESETTING OF HDPS

### A. MSD

At short times, the MSD of reset HDPs starts similarly to that of unperturbed HDPs, as shown in Fig. 5, namely following the power law

**FIG. 5:**
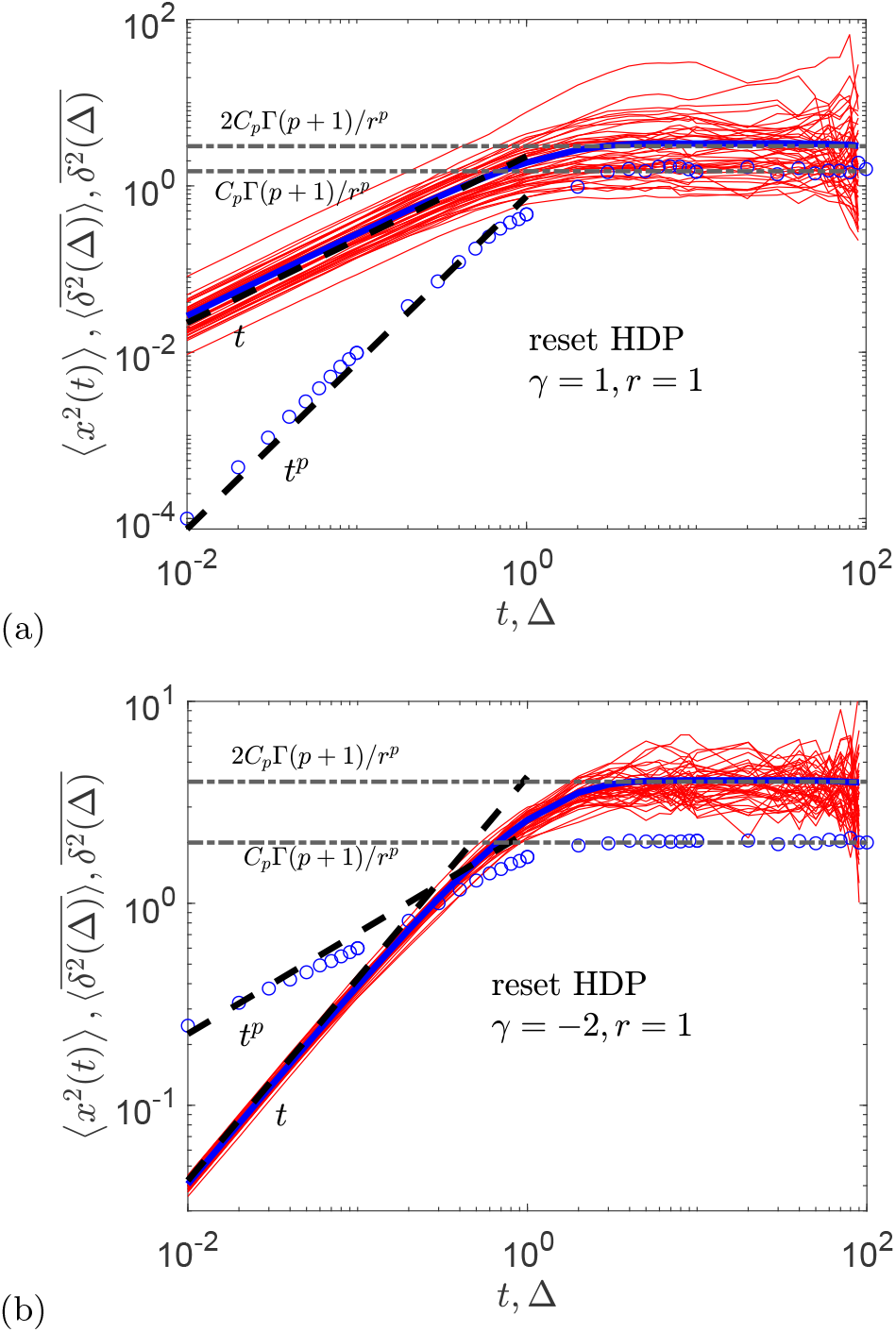
The same as in Fig. 2, but for reset HDPs, computed for the scaling exponents of *D*(*x*) ∼ |*x*| ^*γ*^ being *γ* = 1 (panel (a)) and *γ* = −2 (panel (b)), corresponding to super- and subdiffusive HDPs, respectively. The theoretical short-time asymptotes (37) and (46) and the NESS-related long-time MSD and mean TAMSD plateaus given by expressions (40) and (44), respectively, are the dashed black lines. The magnitude of the diffusivity (17) is fixed in simulations to *D*_0_ = 1.

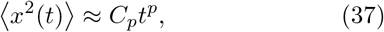

where the scaling exponent of the MSD, *p*(*γ*), is given in terms of the exponent of the space-dependent diffusivity form (17) by^146^

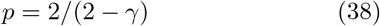

and the prefactor *C*_*p*_ in (37) is^146^

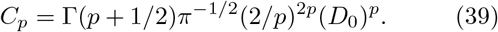

The MSD plateau (see App. B for the derivation)

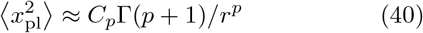

is realized at long times (in the NESS). This expression for the stagnating MSD features the same functional form as the MSD plateau of the reset-FBM process in Eq. (25), with the pair of parameters {*p, C*_*p*_} effectively playing the role of {2*H, K*_2*H*_}. The typical time at which the MSD plateau (40) starts to be followed can be estimated—from equating the growth law (37) and plateau height (40)—as

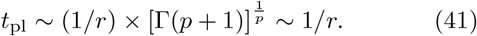

### B. PDF

The approximate PDF of reset HDPs at intermediate-to-long separations follows from the general consideration for reset HDP-FBM processes (see App. C for the derivation, and also Sec. VII) at 2*H* = 1 as

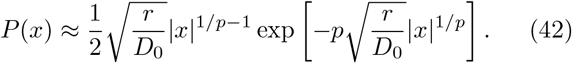

This PDF (truly normalized at *H* = 1*/*2, see Fig. S7) has a Laplacian-like shape in variable |*x*|^1*/p*^ as compared to the Gaussian-like shape of the PDF of nonreset HDPs given by expression (C1). Additionally, for *p* = 1 it turns into the PDF of canonical reset BM^13,14,25,53,54^ given by

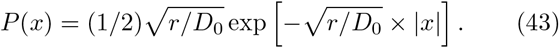

Note that the PDF form (42) cannot predict trimodality observed in computer simulations for reset HDPs with scaling exponents *γ* < 0 because the ideal theoretical consideration yielding the free-HDP PDF (C1) is based upon assuming *infinite* diffusivity at the origin (that, in turn, instantly relocates the particles from there, so that one expects *P* (*x* = 0) = 0), while in the *in-silico-*reality of computer simulations the diffusion coefficient for subdiffusive HDPs has to have finite values at the origin, see Eq. (18), yielding *P* (0) ≠ 0. We quantify the physical reasons of trimodality of PDFs and the dependence of PDF heights at the origin for a general scenario of HDP-FBM processes in Sec. VII B below.

For reset subdiffusive HDPs (*γ* < 0) in the NESS for the trimodal PFD shapes observed in simulations (at certain conditions, see Sec. VII B) the two side peaks stem from the spreading dynamics of nonreset HDPs, while the central peak at *x* = 0 emerges due to resetting to the origin, see Fig. 6. The PDFs for reset superdiffusive HDPs, with the diffusivity exponents 2 > *γ* > 0, have a single peak/cusp at the origin, due to the return events and the influx of particles at *x* = 0, leaving the general PDF shape however largely unaltered.

**FIG. 6:**
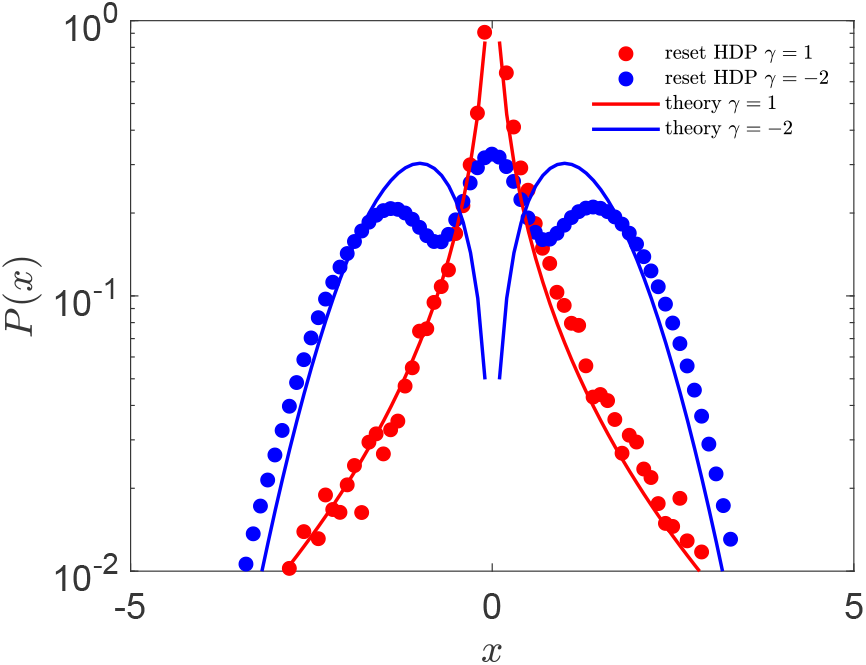
Shapes of the PDFs of reset HDPs plotted for the same scaling exponents of the diffusivity as in Fig. 5, and at diffusion time *T* = 10^2^ in the NESS, as indicated in the legends. The theoretical asymptote (42) is the solid curve.

**FIG. 7:**
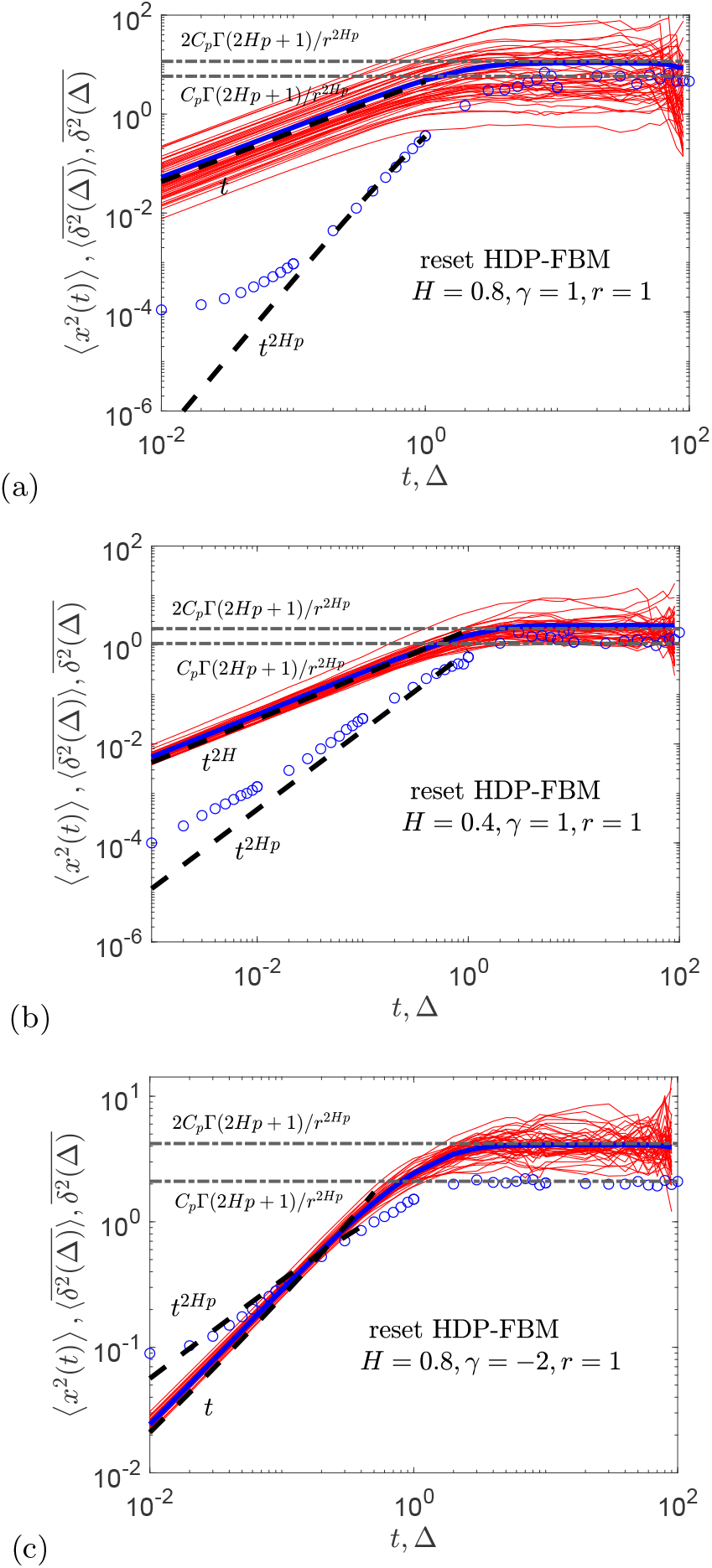
The same as in Fig. 2 but for the generalized reset HDP-FBM process, computed for the parameters as indicated in the plots. The short-time predictions (48) and (55) and the long-time plateaus (48) and (56) for the MSD and mean TAMSD, respectively, are the dashed and dot-dashed lines.

**FIG. 8:**
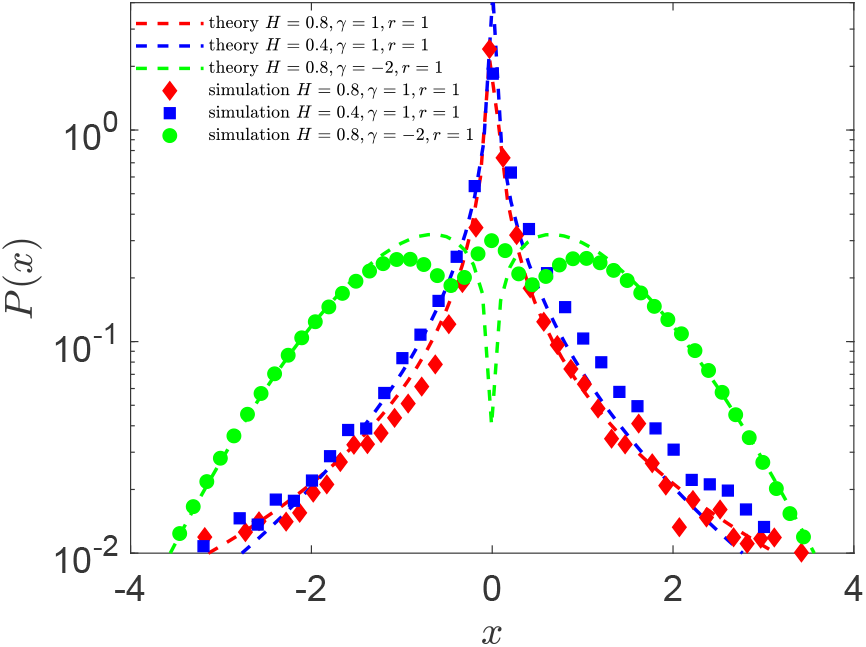
The same as in Fig. 3 but for reset HDP-FBM plotted for diffusion time *t* = *T* = 10^2^ and for the parameters of Fig. 7 (as indicated in the legend).

**FIG. 9:**
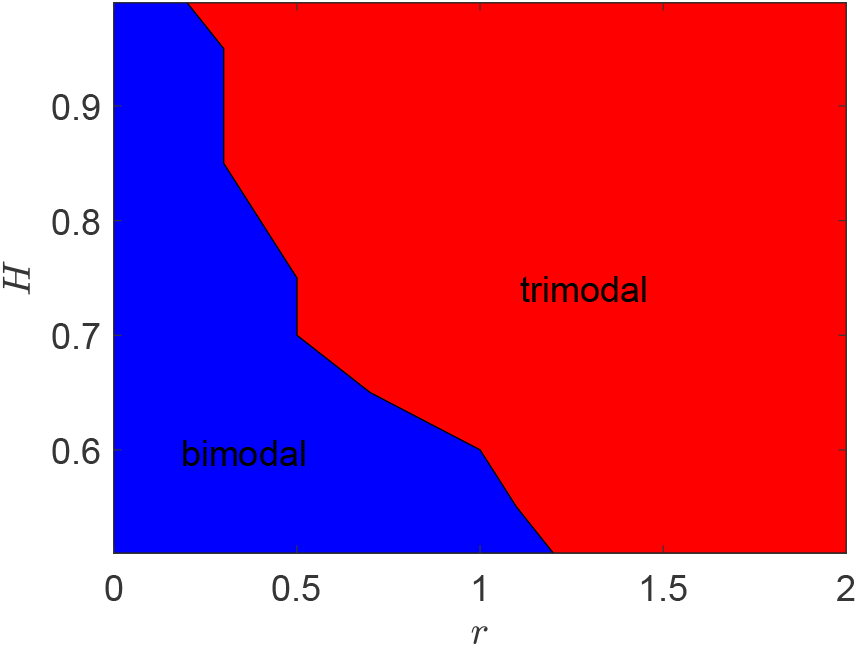
Diagram of bimodal and trimodal PDF shapes for the reset generalized HDP-FBM process in the plane of re-setting rates *r* and Hurst exponents *H*, plotted for the HDP-diffusivity exponent *γ* = −2. The region 0 *< H <* 1*/*2 is not accessible in the simulations of subdiffusive parental HDPs (which are yielding bimodal PDFs being transferred into tri-modal PDFs by resetting). Other parameters: *T* = 10^2^, *dt* = 10^−2^, and *N* = 10^4^.

### C. TAMSD

We start with the long-lag-time behavior here, where a plateau of the mean TAMSD develops, as demonstrated in Fig. 5. The height of this plateau is twice that of the MSD plateau in expression (40), namely

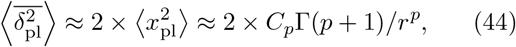

similarly to the TAMSD-versus-MSD plateaus for reset FBM. At short lag times, the linear growth of the mean TAMSD with lag time that is known for HDPs^146,150^,

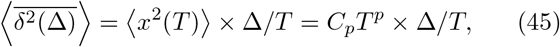

stays unaltered for HDPs in the presence of Poissonian resetting (2), see Fig. 5. Specifically, using the input from computer simulations regarding the linear growth at short lag times and the TAMSD plateau (44) at long lag times, the following approximate evolution of the TAMSD with the lag time can be proposed,

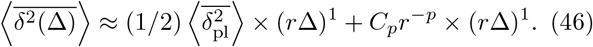

The second term in this expression is analogous to that in Eq. (45), with the inverse reset rate playing the role of the trajectory length in the prefactor, i.e. 1*/r* ↔ *T*.

The spread of individual TAMSD trajectories for subdiffusive reset HDPs with constant-rate resetting becomes relatively small, see Fig. 5b. It is visible in particular for small scaling exponents *γ* when the nonreset HDPs are only weakly nonergodic (with the MSD and mean TAMSD being close in magnitude and in values of their scaling exponents). For superdiffusive HDPs with resetting, the spread of individual TAMSDs is comparatively large, see the behaviors of the MSD and mean TAMSD for *γ* = −2 and *γ* = 1 illustrated in Fig. 5.

### D. EB

The degree of irreproducibility of TAMSD realizations and nonergodicity for reset HDPs depends on the sub-versus superdiffusive nature of nonreset HDPs. In particular for superdiffusive reset HDPs, similarly to the results for superdiffusive reset FBMs in Fig. 4, the EB parameter exhibits a maximum at intermediate rates of reset, see the results shown in Fig. 10 for the case *H* = 1*/*2. For subdiffusive reset HDPs, the EB parameter reveals a plateau in the limit of rare resetting, *r* → 0 (with the height not very sensitive to the exponent *γ*). In the limit of strong resetting, on the other hand, both sub- and superdiffusive HDPs feature the EB parameter decreasing rapidly as ∼ 1*/r* with the rate of exponential resetting, as illustrated in Fig. 10. We thoroughly describe the results for EB variation for a more general process of HDP-FBM in Sec. VII D.

**FIG. 10:**
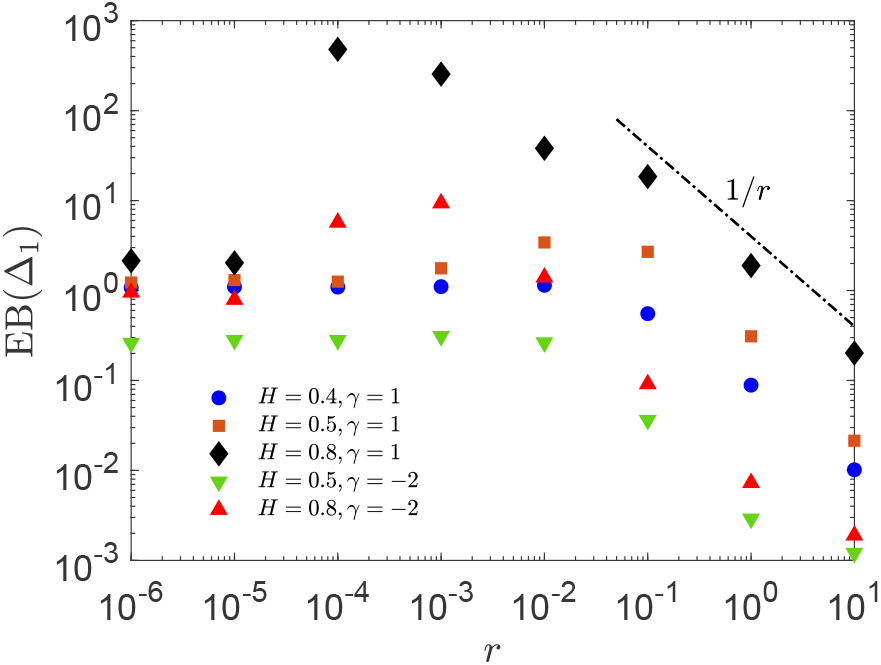
The same as in Fig. 4, but showing the ergodicity breaking parameter for the reset HDP-FBM processes EB(Δ1 = 10^−2^) versus the rate of resetting, for *H* and *γ* exponents as indicated in the legend. The asymptote (35) is the black dot-dashed line shown at intermediate-to-large *r*. Parameters: *T* = 10^2^ and *dt* = 10^−2^.

As a comparison, for interval-confined nonreset HDPs the spread of TAMSD realizations at short lag times was severely restricted by confinement and the values of the EB parameter were shown to decrease as ∼ 1*/T* with the trajectory length *T*, even in the very confined scenarios, see Figs. 4 and 6(b,d) in Ref.^148^. We stress that the MSD-TAMSD inter-relation 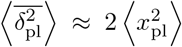 was also valid for the interval-confined HDPs in the limit of long times, after multiple “reflections” of the particles from the confining walls took place.

## VII. RESETTING OF HDP-FBM

### A. MSD

The MSD of reset HDP-FBM at short times starts as for the unperturbed process^150^, following the law

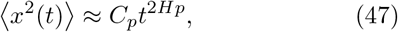

while at long times the NESS plateau of the MSD emerges, with the height (see App. B for the derivation)

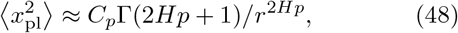

as shown in Fig. 7. This general behavior of the MSD is expected and analogous to that of the parent processes of reset FBM and reset HDPs (considered in Secs. V A and VI A, correspondingly).

### B. PDF

The approximate PDF of a stochastically reset HDP-FBM process at intermediate-to-long distances from the origin, given by expression (see App. C for the derivation)

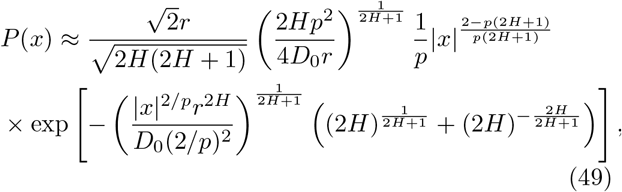

is shown in Fig. 8, revealing a good agreement with the results of our computer simulations.

Trimodal PDFs for reset HDP-FBM processes in the NESS are selected as those shapes having three—rather that one or two—points with zero derivative of the PDF with respect to the coordinate. The necessary condition for this is *γ* < 0 (subdiffusive parental HDPs) and superdiffusive Hurst exponents *H* (as we conclude from the region in the plane of {*H, γ*} amenable for computer simulations, see Fig. 1 in Ref.^150^). The diagram of trimodal PDF shapes in the plane {*H, r*} for a fixed value of the diffusivity exponent *γ* = −2 is presented in Fig. 9. We find that high resetting rates and strongly superdiffusive parental FBMs promote the emergence of PDF trimodality for reset HDP-FBM processes (trimodality is clearly a function of the HDP exponent *γ* < 0 too (the results of simulations for other *γ* values are not shown). In this region of large resetting rates *r* and high Hurst exponents *H* the PDF peak at the reset position *x* = 0 is most distinctly pronounced.

Note that the domain of HDP exponents *γ* < 0 and Hurst exponents *H* < 1*/*2 is not allowed for our specific simulation procedure employed for HDP-FBM processes, see Ref.^150^: thus, we cannot check if trimodal PDFs are present for HDP-FBM also at *H* < 1*/*2 and subdiffusive HDPs (the “forbidden region” of parameters). The region of exponent variation in Figs. S12 and 9 is such that only *γ* < 0 (*p* < 1) and superdiffusive FBMs with *H* > 1*/*2 are allowed (again, see Fig. 1 in Ref.^150^)

Now we rationalize the height of the resetting-induced PDF peak at the origin. Because of regularization of sub-diffusive HDPs via the diffusivity Ansatz (18), the particles returned to the origin start spreading effectively not according to HDP-FBM, but rather as conventional reset FBM. The PDF peak at *x* ≈ 0 cannot be captured by the Laplace-method-based PDF (49) applicable at intermediate-to-large separations from the origin. For larger *r* values, however, the fraction of diffusing particles returned to the origin is comparatively large so that FBM dominates the overall dynamics, while the HDP-based spreading dynamics is not yet established for this process. Therefore, at larger *r* we expect the scaling for the PDF height at the origin, *P* (*x* = 0), for reset HDP-FBM process to follow that of simple reset FBM, namely

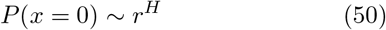

given by (29). This asymptote is indeed found to fit the results of computer simulations rather closely for the regime of frequent resetting, as illustrated in Fig. S12.

For small *r* values the PDF value at *x* = 0 is rather HDP-dynamics dominated, as for a free process of HDP-FBM^150^. Therefore, for rare resetting for reset HDP-FBM we predict the following relation

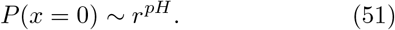

This scaling is reminiscent of that found for free HDP-FBM^150^, with the inverse rate of resetting playing the role of the trajectory length *T*, as intuitively expected,

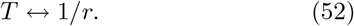

Performing simulations for two different simulation time-steps, in Fig. S12 we show that the differences for the heights of the PDF at the origin do exist, but they are not substantial so that the theoretically predicted asymptotic laws (50) and (51) for *P* (*x* = 0) are still valid.

### C. TAMSD

For reset HDP-FBM for the choice of exponents 0 *< H <* 1*/*2 the leading TAMSD term scales sublinearly as

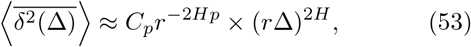

while for 1*/*2 *< H <* 1 the leading term grows linearly,

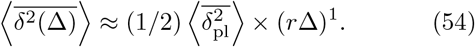

Generally, the mean TAMSD of the reset HDP-FBM process is a combination of these two terms,

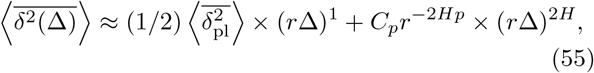

whereas at long lag times the mean-TAMSD plateau is realized with twice the height of the MSD plateau given by (48), i.e.

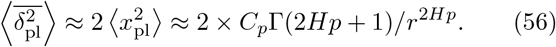

These expressions present a natural continuation of the results for reset FBM and HDPs and they enable excellent quantitative fit of the simulation data for varying model parameters and scaling exponents (*γ* and *H*), see Fig. 7. We emphasize that the sublinear (53) and linear (54) short-lag-time scaling of the TAMSD for reset HDP-FBM are “inherited” from the respective scaling laws for reset FBM, Eqs. (30) and (31), while the the height of the TAMSD plateau in the NESS given by (56) depends on the “intensity” of the HDP-driven dynamics, *C*_*p*_.

Note that for HDPs we generally use slightly nonzero reset positions. In Fig. 7 for *γ* > 0 we set

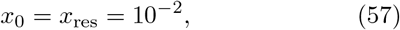

while for *γ <* 0 the reset position was fixed at *x*_0_ = 0.3 (to shorten the transient short-time regime where the theoretical and simulation-based MSD results somewhat differ [due to expected effects of initial-position relaxation^146,147^]).

### D. EB

The spread of individual TAMSDs of stochastically reset HDP-FBM depends—in addition to the leading dependence on resetting rate *r*—on the values of the Hurst exponent *H* and the HDP-diffusivity exponent *γ*. Performing simulations for systematically varying *r* we observe that, especially for superdiffusive FBM being the parent process for HDP-FBM, the dependence of EB on *r* is nonmonotonic. Similarly to the EB(*r*) dependence for reset FBM, the variation of EB(*r*) for HDP-FBM exhibits a maximum at intermediate rates of resetting. For these conditions, as *r* →0 the EB parameter smoothly goes to the respective values for the nonreset process.

In contrast to FBM, for HDP-FBM processes in the absence of resetting the EB values are not small because the parent HDP process is by itself already nonergodic, with MSD≠TAMSD and finite EB values even for infinitely long trajectories^146,147^ (the general features of EB(*p*) variation for HDPs are similar to those of CTRWs^147^). Therefore, for the trajectories of a finite length and at Δ*/T* ≪1 the variations in the magnitudes of short-lag-time TAMSD realizations—characterizing different TAMSD-based diffusivities or transport coefficients—are distinctly visible.

In the opposite limit of very frequent resetting, again similarly to the EB parameter of reset FBM, a power-law decay EB(*r*, Δ_1_) ∼1*/r* is detected in simulations, see Fig. 10. This decay of EB(Δ_1_) at high rates of resetting is universal, being detected for all choices of exponents *H* and *γ* of reset HDP-FBM processes, as well as for trajectories of different lengths, see Fig. S13. The detailed analysis of positioning of this EB(*r*)-maximum as a function of exponents *H* and *γ* as well as of trajectory length *T* is beyond the scope of the current study.

The analysis of whether—for a fixed FBM exponent *H* and resetting rate *r*—the spread of TAMSDs increases as the exponent *γ* deviates from the “most ergodic” value *γ* = 0 [expected to yield smallest EB values] towards negative *γ* for subdiffusive and positive *γ* for superdiffusive HDPs can also be performed. Its results can then be compared to the theoretical predictions for the EB(*γ*)-dependence for nonreset HDPs^146^. All these issues deserve a special theoretical consideration [especially if they turn to be of interest for the experimental resetting community].

**TABLE I:**
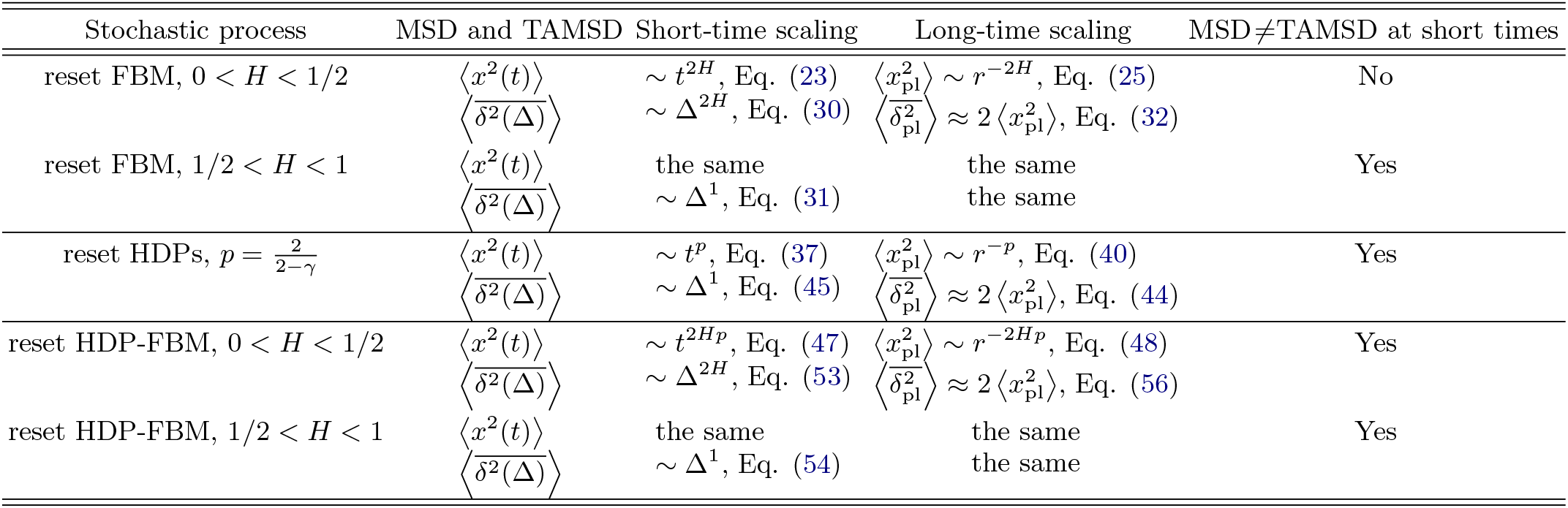
Collection of the main asymptotic results for the MSD and mean TAMSD of reset FBM, HDPs, and HDP-FBM, both at short and long times. The conclusions regarding the nonequivalence of both averages (at short times) are listed in the last column.

## VIII. DISCUSSION AND CONCLUSIONS

The current study is a “Pitot drop”^213^ to a “tsunami” of recent resetting-related publications. This small contribution contains, however, vital single-trajectory-based concepts of the TAMSD and the distribution of TAMSDs ubiquitously used in the analysis of time series from numerous single-particle-tracking experiments. These concepts will hopefully be useful and productive for theoretical studies of other reset stochastic processes (see Sec. VIII C) as well as for experimental resetting setups.

### A. Summary of the main results

The constant-rate Poissonian-resetting setup has been employed to the initially ergodic long-time-memory process of FBM and the initially nonergodic but Markovian HDPs in order to study certain resetting effects onto the ensemble- and time-averaged characteristics of the particle-spreading dynamics (in terms of the MSD and TAMSD, the PDF, and the ergodicity breaking parameter EB). These two widely used processes very different in their stochastic nature, as well as the “compound process”, exemplify how resetting events can smear out the initial distinctions between FBM and HDPs yielding often similar behaviors for many standard quantifiers, as summarized in Tab. I.

#### 1. Reset FBM

We detected nonequivalence of the MSD and mean TAMSD for reset superdiffusive and the equivalence for subdiffusive reset FBM. Specifically, we found that both for sub- and superdiffusive FBM processes the MSD starts (as for free FBM) with ∼ *t*^2*H*^ scaling, while the TAMSD starts sublinearly in lag time for subdiffusive and linearly for superdiffusive reset FBM. In the long-time limit, both the MSD and mean TAMSD revealed the plateau-like behaviors, with the height of the TAMSD in this “stagnating” regime being twice that of the MSD, 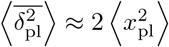, see also Tab. I.

Regarding the PDF of reset FBM in the NESS, we quantified in simulations and described analytically the shapes both at intermediate-to-large separations from the origin, as well as the height of the PDF at the origin (representative of the relative fraction of particles returned at the initial position by the restart events).

Depending on the resetting rate *r*, more frequent re-setting was shown to be capable of both impeding and enhancing the degree of spreading of the magnitudes of individual TAMSDs of reset FBM. The nonmonotonic behavior of EB versus *r* and resetting-induced nonergodicity we discovered in simulations was most pronounced for highly superdiffusive reset FBM, at *H* →1. In the strong-resetting limit, we found a simple power-law decrease EB(*r*) ∼ 1*/r* for reset superdiffusive FBM that can be checked/probed experimentally.

#### 2. Reset HDPs

For reset HDPs, at short times we observed the MSD growing unperturbed as ∼ *t*^*p*^ and the mean TAMSD growing linearly with the lag time, the same scaling relations as for nonreset HDPs^146,147,150^, and thus WEB and MSD-TAMSD nonequivalence is observed for reset HDPs (see also Tab. I). Similarly to reset FBM, for reset HDPs we found that the long-time plateau of the mean TAMSD in the NESS is twice as high as the MSD plateau, as we quantified analytically in excellent agreement with the *in-silico* findings. The relation 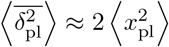 valid for all the reset processes studied here is our **1st key result**.

#### 3. Reset HDP-FBM

For the generalized process of reset HDP-FBM we found that upon Poissonian resetting the MSD at short times always starts as ∼*t*^2*Hp*^, while the mean TAMSD starts linearly for the super- and sublinearly ∼ Δ^2*H*^ for the superdiffusive FBM component of reset HDP-FBM. These asymptotes as well as the long-time plateaus with 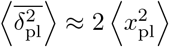 observed in computer simulations agree excellently with the theoretical predictions found. Apart from reset subdiffusive FBM that remained ergodic upon resetting, other pure and “combined” reset processes we considered here have revealed the nonequivalence of the MSD and mean TAMSD at short (lag) times, see Tab. I. This omnipresent MSD-TAMSD nonequivalence at short times upon resetting [even for initially ergodic processes] is our **2nd key result**.

The shape of the PDF computed analytically with the Laplace method was demonstrated to agree well with the numerical results of our Langevin-equation-based stochastic simulations. For subdiffusive HDPs and superdiffusive FBM contributing to the compound (subdiffusive HDP)-(superdiffusive FBM) reset process we detected a novel class of trimodal shapes of the PDF. We unveiled the domain of existence of these trimodal PDFs, with the general conclusion that more frequent resetting and more superdiffusive Hurst exponents of FBM favor trimodal PDF profiles of reset HDP-FBM. We quantified the scaling relations for the peak of the PDF at the origin [the fraction of particles remaining at the reset position], both theoretically and via computer simulations.

Note that although trimodal PDF profiles were considered for specific setups of noninstantaneous resetting for the normal dynamics with certain position-dependent functional forms of the reset speed^45^, the trimodal PDFs we unveiled for reset (subdiffusive HDP)-(superdiffusive FBM) processes are new and universal, being based on the underlying dynamics of the “source” processes of sub-diffusive HDPs with a bimodal PDF and superdiffusive FBM. This emerging PDF trimodality in the NESS for reset subdiffusive HDPs and respective HDP-FBM is our 3rd key result. Such trimodal PDFs can be realizable in diffusion-resetting protocols, even with instant returning of particles to the restart position.

For the EB parameter, similarly to that of reset FBM, at high rates of resetting we discovered a universal power-law decrease of EB of the form EB(*r*) ∼ 1*/r*, with the exact values of EB being sensitive to the values of dynamics-governing exponents *H* and *γ*. At small resetting rates the EB values approach those of the nonreset process, as expected, while a pronounced maximum of EB(*r*) and, thus, resetting-induced nonergodicity emerges at intermediate rates of resetting. This universal nonmonotonic EB(*r*)-dependence—with a prominent maximum followed by EB(*r*) ∝ 1*/r*-decay at large *r*—is our **4**^**th**^ **key result**.

We conclude here stating that the behaviors of the MSD and mean TAMSD in the NESS for FBM, HDPs, and HDP-FBM processes under exponential resetting are functionally *remarkably similar*, alike the found effect of the nonmonotonic variation of EB versus resetting rate. High-rate exponential resetting (naturally) *smears out* the distinctions between initially very different processes we studied here, yielding in the long-time limit similar— possibly, also for other processes under such resetting— functional dependencies for the TAMSD and EB.

### B. Possible effects of initial conditions

We employed the standard [and experimentally relevant] concept of ergodicity as the equivalence of the MSD and mean TAMSD at short times and quantified the spread of individual TAMSDs at Δ = Δ_1_ = *dt* in terms of EB. The initial particle positions were always kept fixed. These conditions are realizable experimentally, with the MSD being computed without subtracting this [nearly-zero] initial position, *x*_0_. From the theoretical perspective, however, considering initial positions of particles being *distributed* according to the long-time PDF in the NESS, *P* (*x*_0_), creates another statistical ensemble and thus offers a different approach to compute averages. The ensemble- and *x*_0_-averaged MSD, computed after this additional averaging as 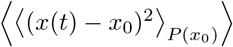, can be a more *theoretically rigorous* measure for assessing the degree of nonergodicity and possible MSD-TAMSD nonequivalence.

We refer the reader to the study^214^ regarding the impact of the starting positions being fixed versus being distributed with the equilibrium PDF on the properties of diffusion in a parabolic potential, the so-called Ornstein-Uhlenbeck (OU) process. It was found, *inter alia*, that the MSD-TAMSD nonequivalence at short lag times indeed *disappears* when the MSD involves second averaging over *P*_eq_(*x*_0_) of all possible starting positions being sampled from the equilibrium-state PDF,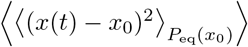. The “confining” OU process is similar to resetting-based protocols also in terms of long-time MSD/TAMSD plateaus featuring 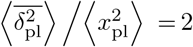 [again, for the starting positions distributed with *P*_eq_(*x*_0_)]^214^. The effects of distributed *x*_0_-positions onto nonergodicity of various reset processes require a separate investigation in the future.

### C. Applications and further developments

Our TAMSD-based approach to reset stochastic processes seems more natural for examining the continuous trajectories, emerging e.g. as outputs in SPT experiments, as compared to the MSD-based methods. Some methods for the analysis of abrupt transitions and the detection of change-points in time series, i.a. in the presence of measurement uncertainties, were developed recently^215^ and applied to predicting the moments of several economic crashes. With our quantifiers, the predicted differences in the short-lag-time scaling of the mean TAMSD for reset FBM—namely, a sublinear and linear TAMSD growth depending on the value of Hurst exponents, see Sec. V C—could help us to assess (e.g., upon varying *H* and comparing with the MSD growth) if the underlying dynamics is indeed of FBM-type.

Straightforward future developments of this study is to apply the TAMSD- and EB-based formulation

- to other stochastic processes with a resetting dynamics imposed—such as CTRWs^16,37,38,55,62,71,124,216^, Lévy walks/flights^217^, SBM^53,54,125^, exponential SBM^131^, diffusion with multiple mobility states, the OU process^214,218^, geometric BM^219–223^, diffusion models with distributed^224,225^ and “diffusing diffusivity” (DD)^144,226–231^ as well as various “hybrid” processes (SBM-HDPs^126^, FBM-DD^144^, SBM-DD^129^, etc.),
- to employ other types of resetting protocols/setups (periodic, power-law, and other functional forms for *ψ*(*r*) distributions; resetting when particular *x*_max_ values are reached, resetting to distributed resetting points, with memory effects, etc.),
- to consider the underdamped versions^129,131^ of anomalous-diffusion processes [with the initially ballistic MSD] with resetting in order to unveil the differences from and the similarities to the behaviors found here for reset FBM, HDPs, and HDP-FBM as well as to mimic relevant experimental situations.

Recently, an experimental realization of diffusion of a colloidal particle with resetting implemented via holographic optical tweezers was reported^67^, possibly allowing position- or energy-dependent resetting of trap-confined beads^36^ and dragging particles^73^ in external traps. The ability and sensitivity of such optical-trap-based setups to infer the underlying stochastic process governing the particles’ motion—as a function of resetting conditions and reset rates being imposed as well as other relevant parameters—remains to be quantified. Moreover, performing such optical-traps experiments with micron-sized beads in crowded environments of living cells—in order to infer whether FBM or viscoelastic diffusion or restricted/compartmentalized diffusion is at play—can potentially have additional complications.

We hope that the theoretical and experimental resetting communities will find our current time-averaging single-trajectory-based approach—with the concepts of TAMSDs and TAMSD-irreproducibility embodied by the EB parameter—useful to infer the degree of nonergodicity for other diffusion models and real physical systems in the presence of [stochastic] resetting dynamics.

## Acknowledgments

A. G. C. gratefully acknowledges the Humboldt University of Berlin for hospitality and support. R. M. thanks the Foundation for Polish Science (Fundacja na rzecz Nauki Polskiej) for support within an Alexander von Humboldt Polish Honorary Research Scholarship.

## Abbreviations

BM: Brownian motion
SBM: scaled BM
FBM: fractional BM
HDPs: heterogeneous diffusion processes
CTRWs: continuous-time random walks
DD model: diffusing-diffusivity model
OU: Ornstein-Uhlenbeck
PDF: probability density function
MSD: mean-squared displacement
TAMSD: time-averaged MSD
WEB: weak ergodicity breaking
NESS: nonequilibrium stationary state
MFPT: mean first-passage time

## Appendix A: Derivation of the mean TAMSD of reset FBM: the limit of short lag times

We start with the autocorrelation function of FBM with Poissonian resetting at a constant rate *r* obtained in Ref.^42^, namely

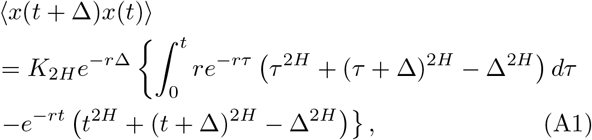

and with the expression for the MSD of SBM under a constant-rate resetting (substituting the SBM exponent *α* via the Hurst exponent *H* of FBM as *α* = 2*H*),^53,54^

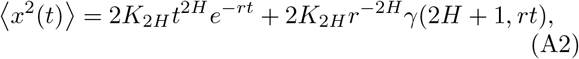

where the (lower) incomplete Gamma function is

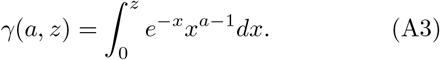

The TAMSD (9), with the integrand

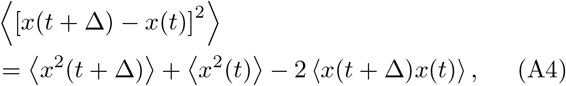

can then be presented as a combination of two terms

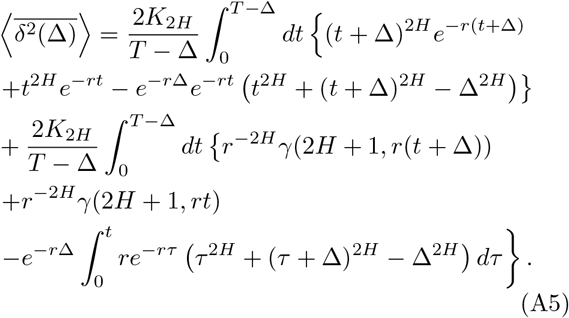

In the limit of vanishing lag times, at Δ*/T* ≪ 1, at the condition of multiple resetting events within a trajectory of length *T*, given by

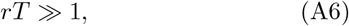

the first integral in Eq. (A5) can be neglected compared to the second term. The second term (after taking the integrals) yields the approximate result for the TAMSD

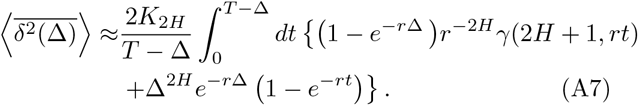

In the limit of long enough trajectories and short enough lag times, at

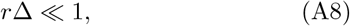

the first term in the integrand of Eq. (A7) yields that for the range of exponents

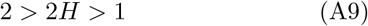

the leading TAMSD behavior is given by

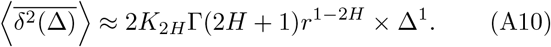

On the contrary, for

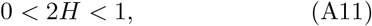

the second term in Eq. (A7) gives the leading contribution to the short-lag-time TAMSD asymptotic that is sublinear in Δ, namely

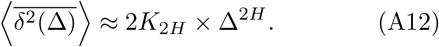

The asymptotes (A10) and (A12)—which are the asymptotic relations (30) and (31) in the main text—can clearly be obtained also via first expanding the integrand of the TAMSD in expression (A5) for small Δ and then taking the limit of long times (not shown in detail).

## Appendix B: MSD of reset HDP-FBM

For the resetting dynamics of a stochastic process with the Gaussian PDF for particle displacements *P*_0_(*x, t*) in the presence of exponential resetting (a constant rate of resetting, Eq. (2)), the PDF of the reset process *P* (*x, t*) can be expressed via the PDF of the unperturbed process *P*_0_(*x, t*) through the following relation^53,54^

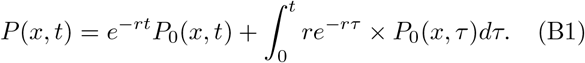

The first term in (B1) signifies the probability of no resetting events taking place from time 0 to time *t* (with the exponentially decaying probability of such an event), while the second term integrates over all multiple resetting events possible to occur in incremental time-steps during the same time period. Multiplying both sides of Eq. (B1) by *x*^2^ and integrating over all possible particle positions one gets for the MSD of the reset HDP-FBM process that

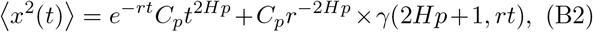

where the definitions/notations (39) and (A3) are used. The height of the MSD plateau at long times (for *rt* ≫ 1 and many resetting events taking place by time *t* [longtime limit of NESS]) is dominated by the second term in Eq. (B2), namely

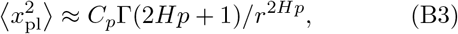

that yields expression (48) in the main text. At short times (when the condition *rt* ≪ 1 is satisfied) the Taylor expansion of (B2) yields that the MSD starts nearly unperturbed, as^150^

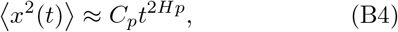

that is expressions (37) and (47) in the main text.

## Appendix C: Derivation of the PDF for reset HDP-FBM processes

Starting with the PDF of HDP-FBM in the absence of resetting^150^,

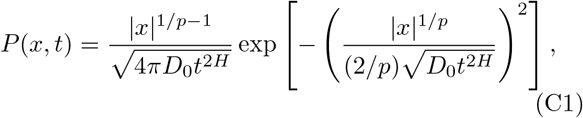

and using the relation (B1) connecting the standard/nonreset and the reset PDFs, in the limit of long times (when the term of no-resetting up to time *t, e*^−*rt*^*P*_0_(*x, t*) in (B1), can be neglected) we arrive at

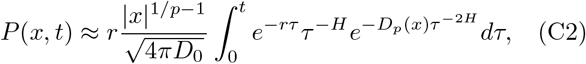

where we defined for brevity

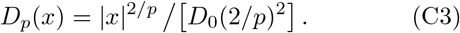

Following the strategy outlined in Ref.^53,54^ for the PDF of reset SBM, we use the Laplace method to approximate the exponent-containing integral in (C2). The maximum of a negative-power exponent in (C2) is achieved at the minimum of its argument, namely at

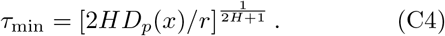

This yield for the second derivative

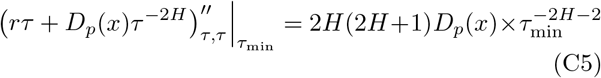

so that the power of the exponent becomes

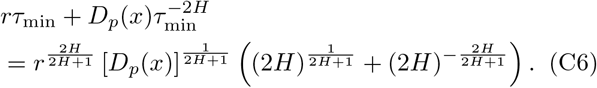

For the prefactor of the exponent in the resulting PDF, 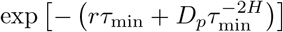, obtained after applying the Laplace method to expression (C2), one gets

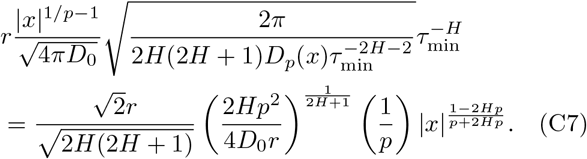

Combining (C6) with (C7) and substituting the explicit *x*-dependence of *D*_*p*_ from (C3), we arrive at the final approximate result for the PDF of HDP-FBM in Eq. (49) of the main text,

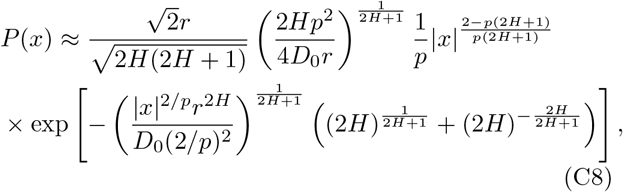

Naturally, in the absence of space-dependent diffusion (when the MSD is linear, *p* = 1) the NESS PDF of HDP-FBM (49) turns after putting 2*H* = *α* into the PDF of SBM with fully renewal resetting^53,54^, see also Eq. (26).

## Appendix D: Supplementary figures

Here we present some auxiliary figures supporting the claims in the main text.

---

**FIG. S1:**
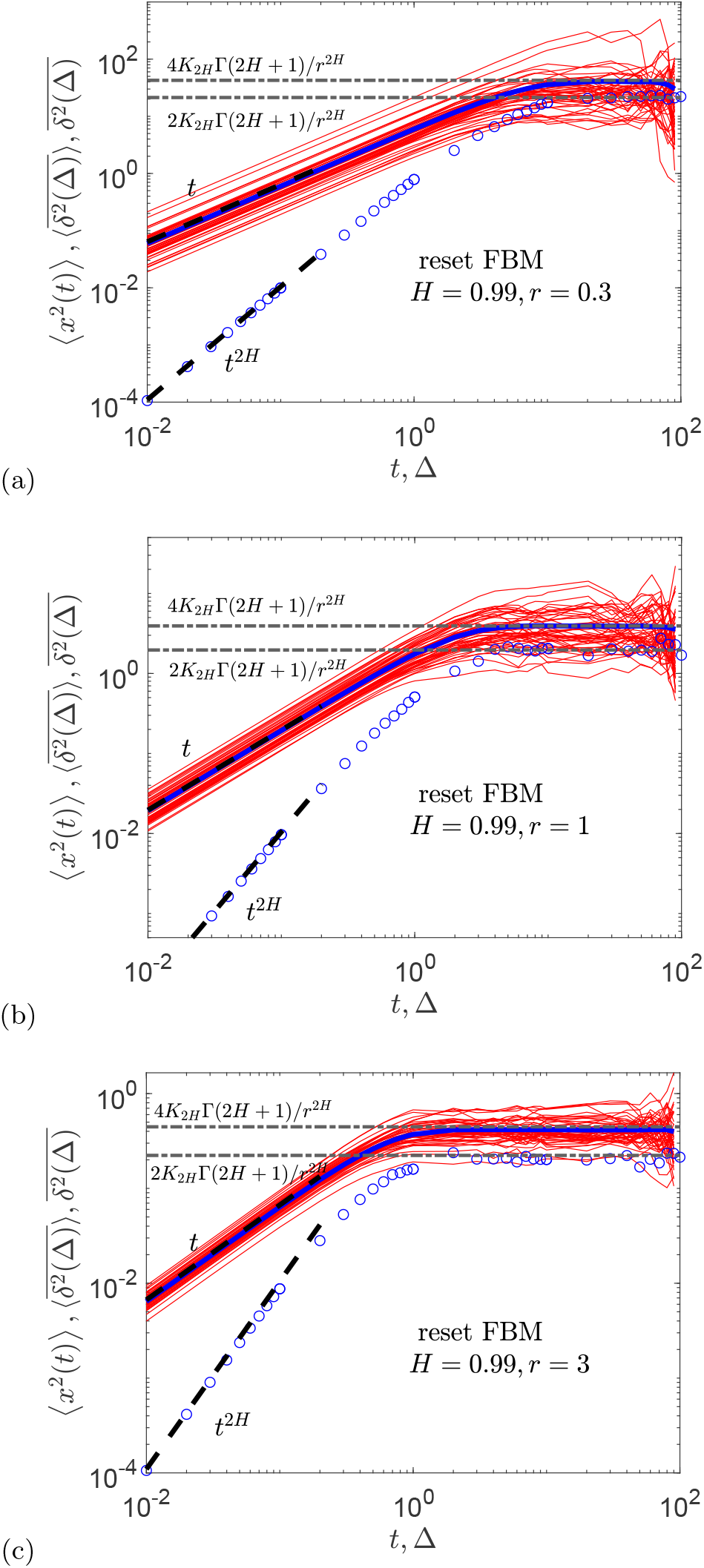
The same as in Fig. 2, for the same parameters, except for *H* = 0.99 and varying resetting rate *r* (the values are indicated in the plots), with the same asymptotes shown.

**FIG. S2:**
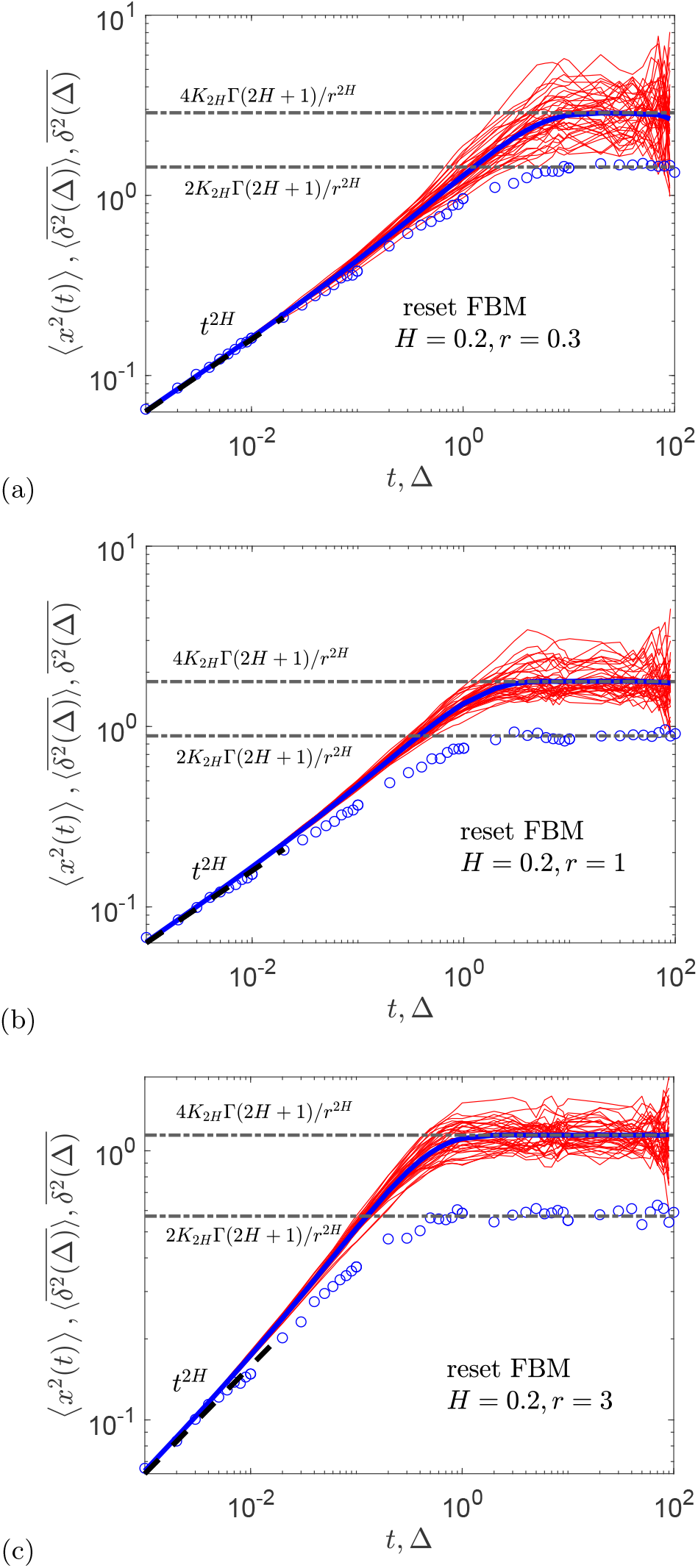
The same as in Fig. S1, for the same parameters except for *H* = 0.2 and *dt* = 10^−3^, computed for varying rate of resetting *r*.

**FIG. S3:**
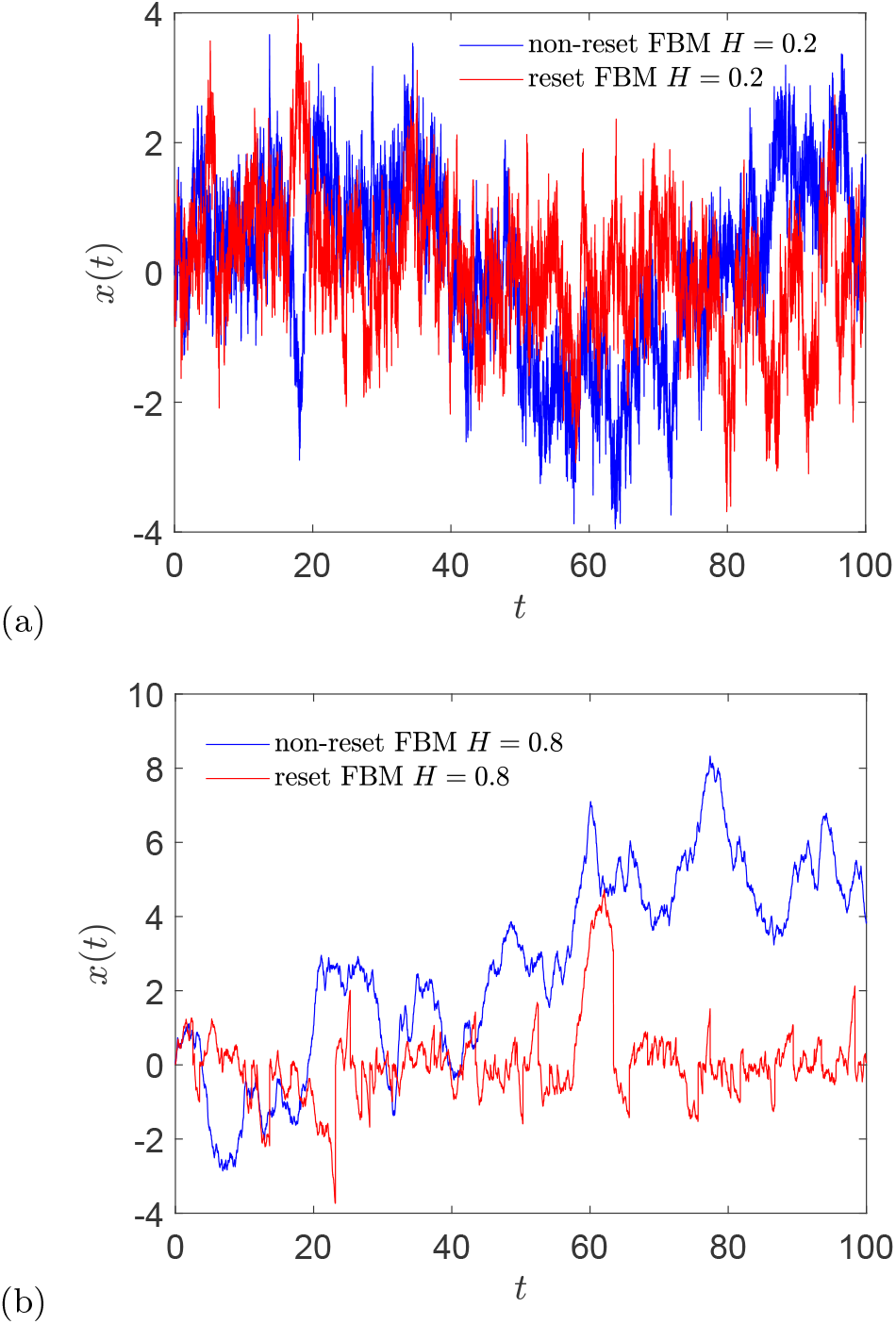
Exemplary trajectories of reset (red) and nonreset (blue) FBM for subdiffusive and superdiffusive Hurst exponents, *H* = 0.2 for panel (a) and *H* = 0.8 for panel (b), with other parameters being the same as in Fig. 2.

**FIG. S4:**
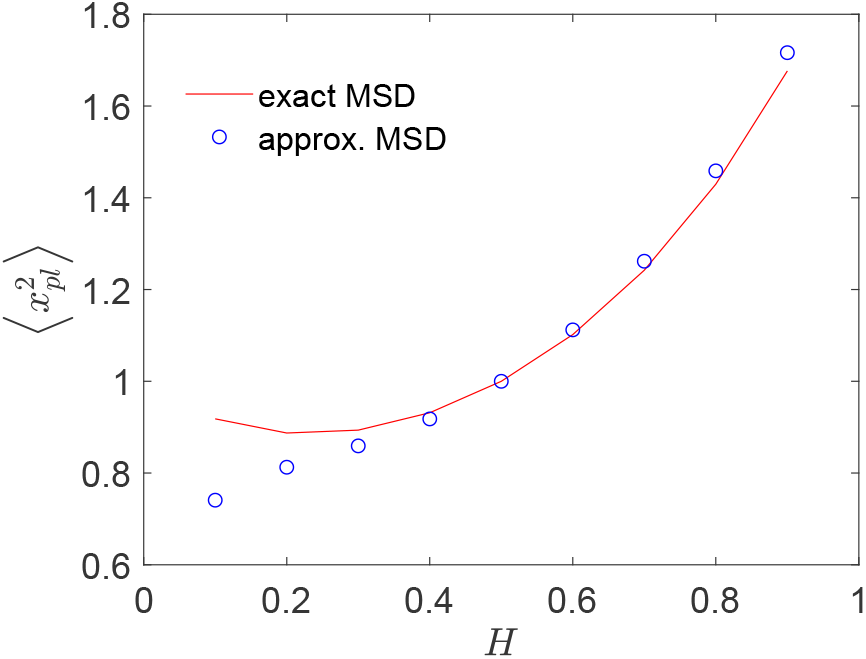
Comparison of the MSD plateaus of reset FBM in the NESS between Eq. (25) and using the approximate PDF form (26), plotted for varying Hurst exponents.

**FIG. S5:**
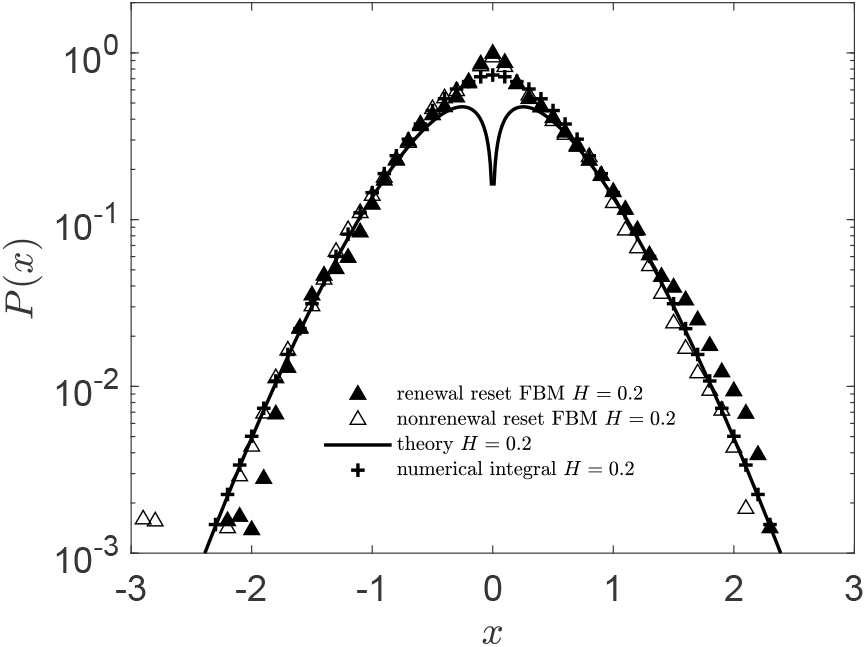
The same as in Fig. 3, for *r* = 10, *dt* = 10^−2^, and *H* = 0.2. The results of numerical integration of the exact PDF expression (B1) are the cross symbols

**FIG. S6:**
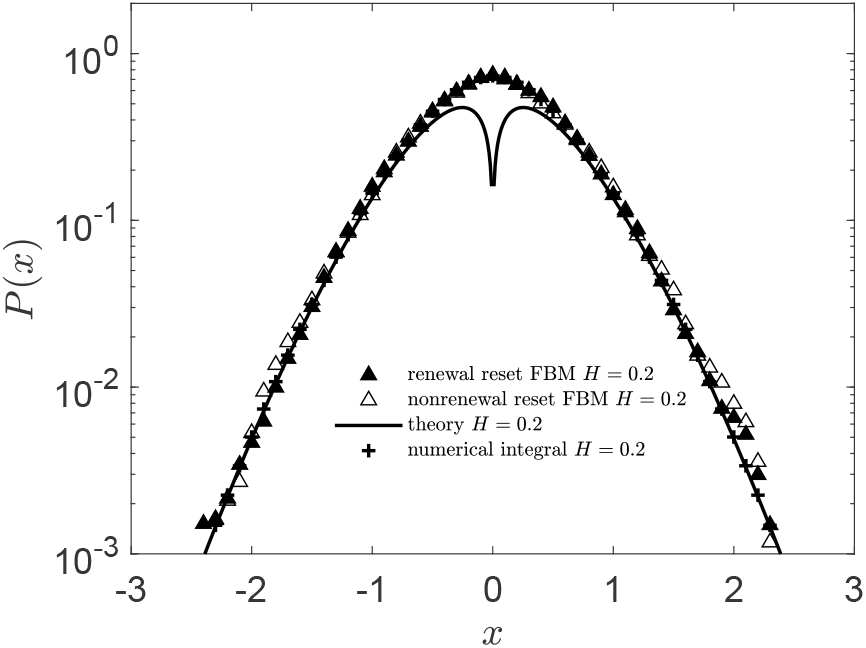
The same as in Fig. S5, for the same parameters, except for *dt* = 10^−3^ and *r* = 10 being used in simulations.

**FIG. S7:**
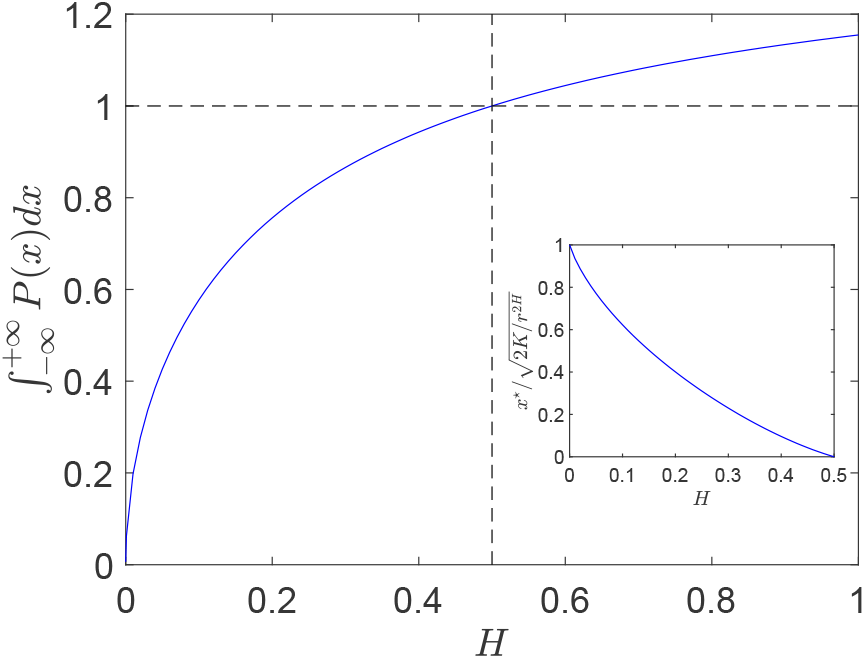
Variation of the integral/norm of the approximate Laplace-method PDF given by (26) computed numerically versus the Hurst exponent *H*. The inset shows the normalized separation *x*^*\**^ expressed by (28) versus *H*.

**FIG. S8:**
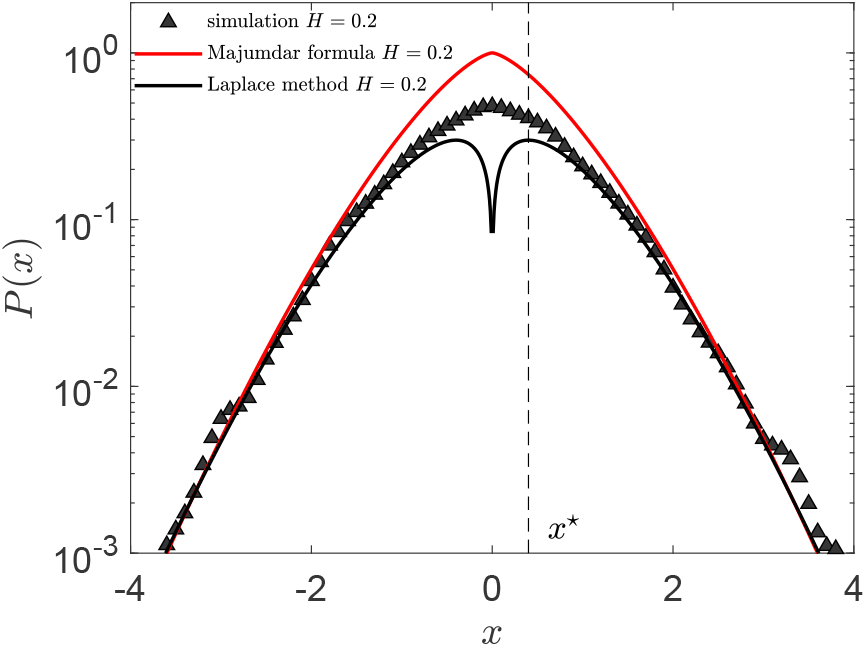
Results of computer simulations of subdiffusive reset FBM compared to the PDF expression (26) and to the leading decay law (27) [derived first in App. 1 of Ref.^25^]. The threshold separation (28) is indicated as the dotted line. Parameters: *H* = 0.2 and *r* = 1.

**FIG. S9:**
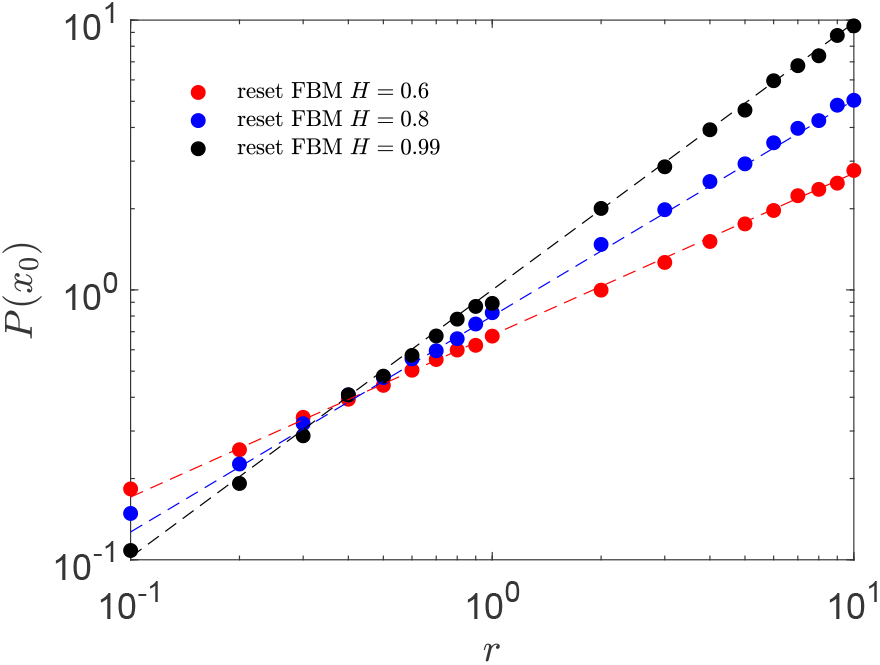
PDF values of reset FBM at *x* = 0 shown for varying reset rates and for different Hurst exponents. The theoretical predictions (29) are the dashed lines of the respective color. Parameters: *T* = 10^2^ and *dt* = 10^−2^.

**FIG. S10:**
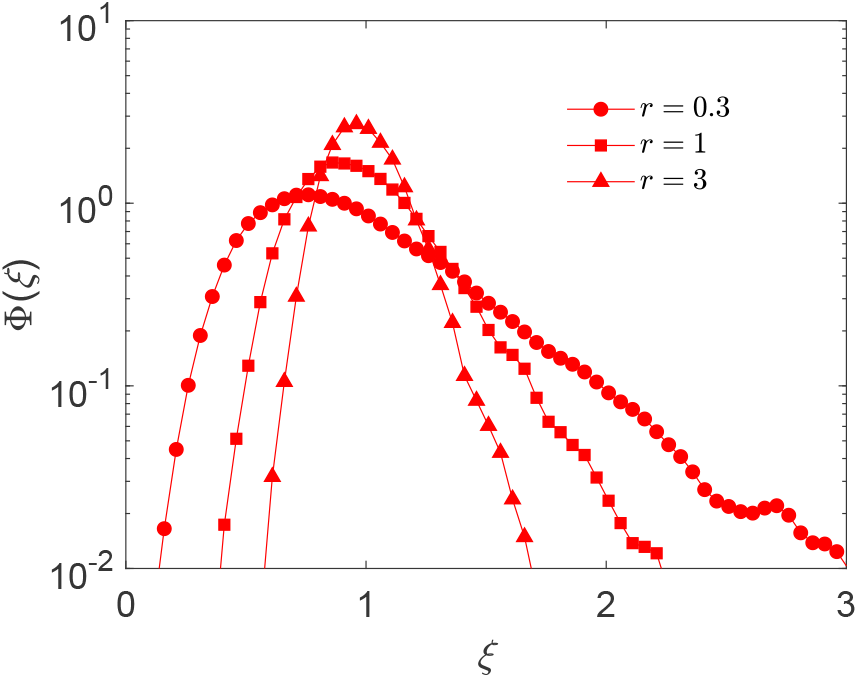
Scatter distribution of TAMSD amplitudes, defined as *ϕ*(*ξ*(Δ)) in Eq. (12), computed at the shortest lag time Δ = Δ1 = 10^−2^ for reset superdiffusive FBM with *H* = 0.8 and for varying resetting rates, for *r* = 0.3, 1, 3, with other parameters being the same as in Fig. 2c.

**FIG. S11:**
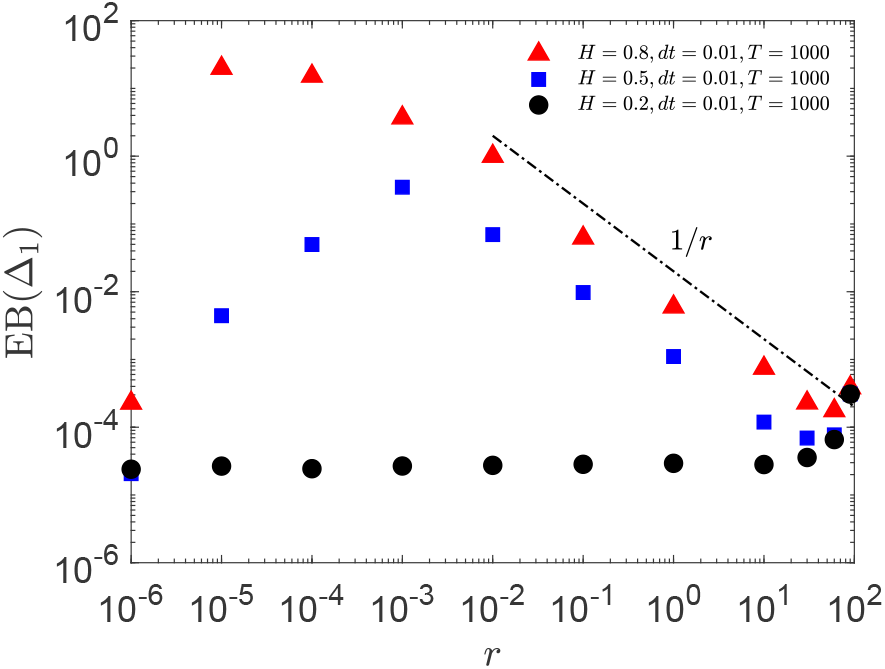
The same as in Fig. 4, computed for the same parameters, except for *T* = 10^3^ (see also the legend).

**FIG. S12:**
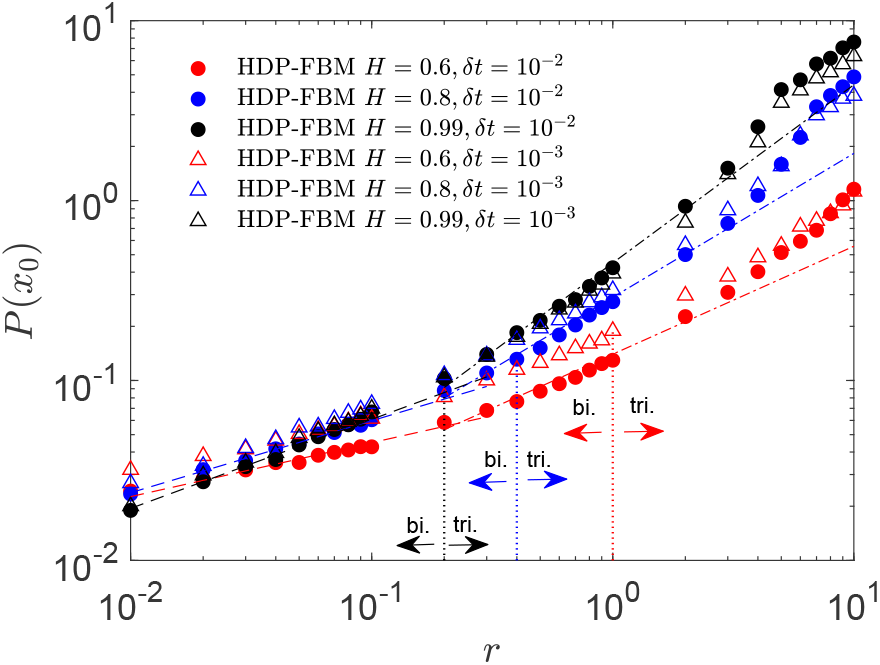
PDF values of the reset HDP-FBM process for *p* = 1*/*2 (or *γ* = −2), with the regions of bimodal (rate resetting) and trimodal (frequent resetting) PDF shapes being indicated for each choice of the Hurst exponent, computed at *x* = *x*_0_ = 0.01 from computer simulations. The analytical asymptotes (29) and (51) are the dashed lines of the corresponding color shown, respectively, in the regime of large and small rates of resetting. The simulation data for *dt* = 10^−2^ and 10^−3^ are shown by filled and empty symbols, correspondingly. Parameters: *T* = 10 with *dt* = 10^−3^ and *T* = 10^2^ with *dt* = 10^−2^, see the legend for details, while ensemble averaging executed over *N* = 8000 HDP-FBM trajectories.

**FIG. S13:**
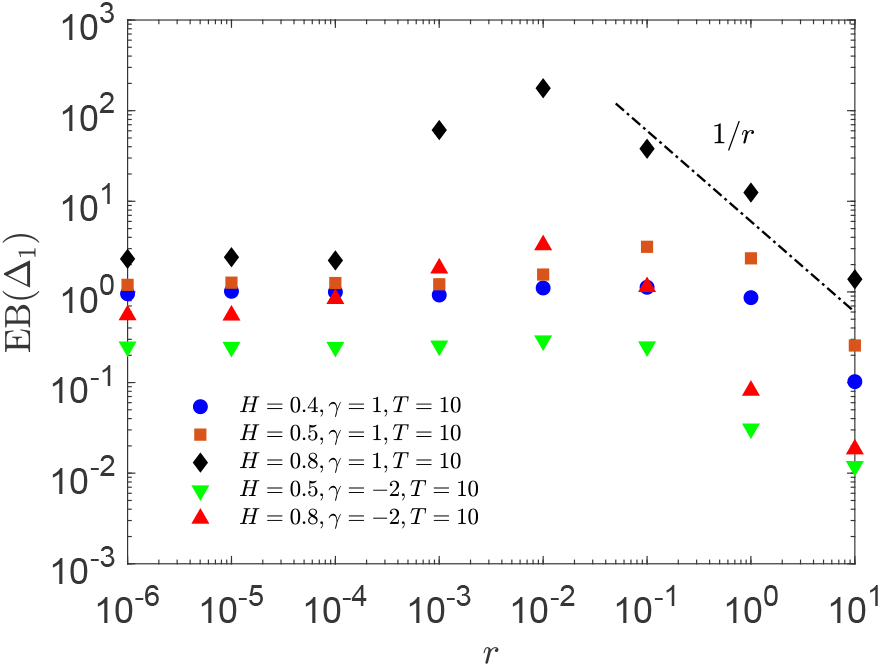
The same as in Fig. 10, for the same parameters, but for *T* = 10.

## References

1 J.-P. Bouchaud and A. Georges, Anomalous diffusion in disordered media: statistical mechanisms, models and physical applications, Phys. Rep. 195, 127 (1990).

2 R. Metzler and J. Klafter, The random walk’s guide to anomalous diffusion: a fractional dynamics approach, Phys. Rep. 339, 1 (2000).

3 S. Burov, J.-H. Jeon, R. Metzler, and E. Barkai, Single particle tracking in systems showing anomalous diffusion: the role of weak ergodicity breaking, Phys. Chem. Chem. Phys. 13, 1800 (2011).

4 I. M. Sokolov, Models of anomalous diffusion in crowded environments, Soft Matter 8, 9043 (2012).

5 E. Barkai, Y. Garini, and R. Metzler, Strange kinetics of single molecules in living cells, Phys. Today 65, 29 (2012).

6 F. Höfling and T. Franosch, Anomalous transport in the crowded world of biological cells, Rep. Prog. Phys. 76, 046602 (2013).

7 Y. Meroz, I. M. Sokolov, and J. Klafter, Test for determining a subdiffusive model in ergodic systems from single trajectories, Phys. Rev. Lett. 110, 090601 (2013).

8 R. Metzler, J.-H. Jeon, A. G. Cherstvy, and E. Barkai, Anomalous diffusion models and their properties: nonstationarity, non-ergodicity, and ageing at the centenary of single particle tracking, Phys. Chem. Chem. Phys. 16, 24128 (2014).

9 Y. Meroz and I. M. Sokolov, A toolbox for determining subdiffusive mechanisms, Phys. Rep. 573, 1 (2015).

10 M. A. F. dos Santos, Analytic approaches of the anomalous diffusion: a review, Chaos, Solitons & Fractals 124, 86 (2019).

11 S. C. Manrubia and D. H. Zanette, Stochastic multiplicative processes with reset events, Phys. Rev. E 59, 4945 (1999).

12 A. Montanari and R. Zecchina, Optimizing searches via rare events, Phys. Rev. Lett. 88, 178701 (2002).

13 M. R. Evans and S. N. Majumdar, Diffusion with stochastic resetting, Phys. Rev. Lett. 106, 160601 (2011).

14 M. R. Evans and S. N. Majumdar, Diffusion with optimal resetting, J. Phys. A 44, 435001 (2011).

15 M. R. Evans, S. N. Majumdar, and K. Mallick, Optimal diffusive search: nonequilibrium resetting versus equilibrium dynamics, J. Phys. A 46, 185001 (2013).

16 M. Montero and J. Villarroel, Monotonic continuous-time random walks with drift and stochastic reset events, Phys. Rev. E 87, 012116 (2013).

17 S. Gupta, S. N. Majumdar, and G. Schehr, Fluctuating interfaces subject to stochastic resetting, Phys. Rev. Lett. 112, 220601 (2014).

18 M. R. Evans and S. N. Majumdar, Diffusion with resetting in arbitrary spatial dimension, J. Phys. A 47, 285001 (2014).

19 L. Kusmierz, S. N. Majumdar, S. Sabhapandit, and G. Schehr, First order transition for the optimal search time of Lévy flights with resetting, Phys. Rev. Lett. 113, 220602 (2014).

20 D. Boyer and C. Solis-Salas, Random walks with preferential relocations to places visited in the past and their application to biology, Phys. Rev. Lett. 112, 240601 (2014).

21 X. Durang, M. Henkel, and H. Park, The statistical mechanics of the coagulation-diffusion process with a stochastic reset, J. Phys. A 47, 045002 (2014).

22 S. Reuveni, M. Urbakh, and J. Klafter, Role of substrate unbinding in Michaelis-Menten enzymatic reactions, Proc. Natl. Acad. Sci. U. S. A. 111, 4391 (2014).

23 T. Rotbart, S. Reuveni, and M. Urbakh, Michaelis-Menten reaction scheme as a unified approach towards the optimal restart problem, Phys. Rev. E 92, 060101 (2015).

24 J. M. Meylahn, S. Sabhapandit, and H. Touchette, Large deviations for Markov processes with resetting, Phys. Rev. E 92, 062148 (2015).

25 S. N. Majumdar, S. Sabhapandit, and G. Schehr, Dynamical transition in the temporal relaxation of stochastic processes under resetting, Phys. Rev. E 91, 052131 (2015).

26 A. Pal, Diffusion in a potential landscape with stochastic resetting, Phys. Rev. E 91, 012113 (2015).

27 S. N. Majumdar, S. Sabhapandit, and G. Schehr, Random walk with random resetting to the maximum position, Phys. Rev. E 92, 052126 (2015).

28 C. Christou, and A. Schadschneider, Diffusion with resetting in bounded domains, J. Phys. A 48, 285003 (2015).

29 S. Eule and J. J. Metzger, Non-equilibrium steady states of stochastic processes with intermittent resetting, New J. Phys. 18, 033006 (2016).

30 S. Reuveni, Optimal stochastic restart renders fluctuations in first passage times universal, Phys. Rev. Lett. 116, 170601 (2016)

31 A. Nagar and S. Gupta, Diffusion with stochastic resetting at power-law times, Phys. Rev. E 93, 060102(R) (2016).

32 J. Fuchs, S. Goldt, and U. Seifert, Stochastic thermodynamics of resetting, EPL 113, 60009 (2016).

33 E. Roldan, A. Lisica, D. Sanchez-Taltavull, and S. W. Grill, Stochastic resetting in backtrack recovery by RNA Phys. J. B 90, 176 (2017).

34 A. Pal, A. Kundu, and M. R. Evans, Diffusion under time-dependent resetting, J. Phys. A 49, 225001 (2016).

35 A. Pal and S. Reuveni, First passage under restart, Phys. Rev. Lett. 118, 030603 (2017).

36 E. Roldan and S. Gupta, Path-integral formalism for stochastic resetting: exactly solved examples and shortcuts to confinement, Phys. Rev. E 96, 022130 (2017).

37 V. P. Shkilev, Continuous-time random walk under time-dependent resetting, Phys. Rev. E 96, 012126 (2017).

38 M. Montero, A. Maso-Puigdellosas, and J. Villarroel, Continuous-time random walks with reset events, Eur. (2018).

39 R. Falcao and M. R. Evans, Interacting Brownian motion with resetting, J. Stat. Mech. 02320, (2017).

40 M. R. Evans and S. N. Majumdar, Run and tumble parpolymerases, Phys. Rev. E 93, 062411 (2016).

41 A.V. Chechkin and I.M. Sokolov, Random search with resetting: a unified renewal approach, Phys. Rev. Lett. 121, 050601 (2018).

42 S.N. Majumdar and G. Oshanin, Spectral content of fractional Brownian motion with stochastic reset, J. Phys. A 51, 435001 (2018).

43 A. Chatterjee, C. Christou, and A. Schadschneider, Diffusion with resetting inside a circle, Phys. Rev. E 97, 062106 (2018).

44 C. Christou, “Non-Equilibrium Stochastic Models: Random Average Process and Diffusion with Resetting”, PhD Thesis, (Universität zu Köln, 2019).

45 A. Pal, L. Kusmierz, and S. Reuveni, Invariants of motion with stochastic resetting and space-time coupled returns, New J. Phys. 21, 113024 (2019).

46 A. Pal and V. V. Prasad, Landau-like expansion for phase transitions in stochastic resetting, Phys. Rev. Res. 1, 032001 (2019).

47 A. Pal, L. Kusmierz, and S. Reuveni, Time-dependent density of diffusion with stochastic resetting is invariant to return speed, Phys. Rev. E 100, 040101(R) (2019).

48 D. Gupta, Stochastic resetting in underdamped Brownian motion, J. Stat. Mech. 033212, (2019).

49 M. A. F. dos Santos, Fractional Prabhakar derivative in diffusion equation with non-static stochastic resetting, MDPI: Physics 1, 40 (2019).

50 A. Maso-Puigdellosas, D. Campos, and V. Mendez, Transport properties and first-arrival statistics of random motion with stochastic reset times, Phys. Rev. E 99, 012141 (2019).

51 J. Masoliver, Telegraphic processes with stochastic resetting, Phys. Rev. E 99, 012121 (2019).

52 T. Robin, L. Hadany, and M. Urbakh, Random search with resetting as a strategy for optimal pollination, Phys. Rev. E 99, 052119 (2019).

53 A. S. Bodrova, A. V. Chechkin, and I. M. Sokolov, Nonrenewal resetting of scaled Brownian motion, Phys. Rev. E 100, 012119 (2019).

54 A. S. Bodrova, A. V. Chechkin, and I. M. Sokolov, Scaled Brownian motion with renewal resetting, Phys. Rev. E 100, 012120 (2019).

55 A. S. Bodrova and I. M. Sokolov, Continuous-time random walks under power-law resetting, Phys. Rev. E 101, 062117 (2020).

56 A. S. Bodrova and I. M. Sokolov, Brownian motion under noninstantaneous resetting in higher dimensions, Phys. Rev. E 102, 032129 (2020).

57 A. S. Bodrova and I. M. Sokolov, Resetting processes with noninstantaneous return, Phys. Rev. E 101, 052130 (2020).

58 G. Mercado-Vasquez, D. Boyer, S. N Majumdar, and G. Schehr, Intermittent resetting potentials, J. Stat. Mech. 113203, (2020).

59 M. R. Evans, S. N. Majumdar, and G. Schehr, Stochastic resetting and applications, J. Phys. A 53, 193001 (2020).

60 C. A. Plata, D. Gupta, and S. Azaele, Asymmetric stochastic resetting: Modeling catastrophic events, Phys. Rev. E 102, 052116 (2020).

61 S. Ray, Space-dependent diffusion with stochastic resetting: a first-passage study, J. Chem. Phys. 153, 234904 (2020).

62 T. Zhou, P. Xu, and W. Deng, Continuous-time random walks and Lévy walks with stochastic resetting, Phys. Rev. Res. 2, 013103 (2020).

63 A. Pal, L. Kusmierz, and S. Reuveni, Search with home returns provides advantage under high uncertainty, Phys. Rev. Res. 2, 043174 (2020).

64 R. K. Singh, R. Metzler, and T. Sandev, Resetting dynamics in a confining potential, J. Phys. A. 53, 505003 (2020).

65 V. Domazetoski, A. Maso-Puigdellosas, T. Sandev, V. Mendez, A. Iomin, and L. Kocarev, Stochastic resetting on comblike structures, Phys. Rev. Res. 2, 033027 (2020).

66 B. Besga, A. Bovon, A. Petrosyan, S. N. Majumdar, and S. Ciliberto, Optimal mean first-passage time for a Brownian searcher subjected to resetting: experimental and theoretical results, Phys. Rev. Res, 2, 032029(R) (2020).

67 O. Tal-Friedman, A. Pal, A. Sekhon, S. Reuveni, and Y. Roichman, Experimental realization of diffusion with stochastic resetting, J. Phys. Chem. Lett. 11, 7350 (2020).

68 A. F. M. dos Santos, Comb model with non-static stochastic resetting and anomalous diffusion, Fractal Fract. 4, 28 (2020).

69 P. C. Bressloff, Queueing theory of search processes with stochastic resetting, Phys. Rev. E 102, 032109 (2020).

70 A. Miron and S. Reuveni, Diffusion with local resetting and exclusion, Phys. Rev. Res. 3, L012023 (2021).

71 V. Mendez, A. Maso-Puigdellosas, T. Sandev, and D. Campos, Continuous time random walks under Markovian resetting, Phys. Rev. E 103, 022103 (2021).

72 T. Sandev, V. Domazetoski, A. Iomin, and L. Kocarev, Diffusion-advection equations on a comb: resetting and random search, MDPI: Mathematics 9, 221 (2021).

73 D. Gupta, C. A. Plata, A. Kundu, and A. Pal, Stochastic resetting with stochastic returns using external trap, J. Phys. A 54, 025003 (2021).

74 C. Monthus, Large deviations at various levels for run-and-tumble processes with space-dependent velocities and space-dependent switching rates, 2103.08885 (2021).

75 P. Singh and A. Pal, Extremal statistics for stochastic resetting systems, 2102.07111 (2021).

76 F. Mori, S. N. Majumdar, and G. Schehr, Detecting nonequilibrium dynamics via extreme value statistics, 2104.07346 (2021).

77 S. Ciliberto, Experiments in stochastic thermodynamics: short history and perspectives, Phys. Rev. X 7, 021051 (2017).

78 D. W. Stephens and J. R. Krebs, ”Foraging Theory”, vol. 1 (Princeton University Press, 1986).

79 G. M. Viswanathan, S. V. Buldyrev, S. Havlin, M. G. E. da Luz, E. P. Raposo, and H. E. Stanley, Optimizing the success of random searches, Nature 401, 911 (1999).

80 F. Bartumeus and J. Catalan, Optimal search behavior and classic foraging theory, J. Phys. A 42, 434002 (2009).

81 N. E. Humphries, H. Weimerskirch, N. Queiroz, E. J. Southall, and D. W. Sims, Foraging success of biological Lévy flights recorded in situ, Proc. Natl. Acad. Sci. U. S. A. 109, 7169 (2012).

82 T. H. Harris et al., Generalized Lévy walks and the role of chemokines in migration of effector CD8+ 1 T cells, Nature 486, 545 (2012).

83 G. Ariel, A. Rabani, S. Benisty, J. D. Partridge, R. M. Harshey, and A. Be’er, Swarming bacteria migrate by Lévy walk, Nature Comm. 6, 8396 (2015).

84 T. Shokaku, T. Moriyama, H. Murakami, S. Shinohara, N. Manome, and K. Morioka, Development of an automatic turntable-type multiple T-maze device and observation of pill bug behavior, Rev. Sci. Instrum. 91, 104104 (2020).

85 B. Guinard and A. Korman, Intermittent inverse-square Lévy walks are optimal for finding targets of all sizes, Science Adv. 7, eabe8211 (2021).

86 W. Bell, ”The Behavioural Ecology of Finding Resources”, (Springer, Netherlands, 1990).

87 P. Turchin, ”Quantitative Analysis of Movement”, (Sinauer Associates Inc., Sunderland, MA, 1998).

88 G. M. Viswanathan, M. G. E. da Luz, E. P. Raposo, and H.E, Stanley, ”The Physics of Foraging: An Introduction to Random Searches and Biological Encounters”, (Cambridge University Press, 2011).

89 O. Benichou, C. Loverdo, M. Moreau, and R. Voituriez, Intermittent search strategies, Rev. Mod. Phys. 83, 81 (2011).

90 O. Vilk, Y. Orchan, M. Charter, N. Ganot, S. Toledo, R. Nathan, and M. Assaf, Ergodicity breaking and lack of a typical waiting time in area-restricted search of avian predators, 2101.11527 (2021).

91 M. Slutsky, M. Kardar, and L. A. Mirny, Diffusion in correlated random potentials, with applications to DNA, Phys. Rev. E. 69, 061903 (2004).

92 M. Slutsky and L. A. Mirny, Kinetics of protein-DNA interaction: facilitated target location in sequence-dependent potential, Biophys. J. 87, 4021 (2004).

93 I. M. Sokolov, R. Metzler, K. Pant, and M. C. Williams, Target search of N sliding proteins on a DNA, Biophys J. 89, 895 (2005).

94 A. G. Cherstvy, A. A. Kolomeisky, and A. A. Kornyshev, Protein-DNA interactions: reaching and recognizing the targets, J. Phys. Chem. B 112, 4741 (2008).

95 A. B. Kolomeisky, Physics of protein-DNA interactions: mechanisms of facilitated target search, Phys. Chem. Chem. Phys. 13, 208 (2011).

96 E. Kussell, and S. Leibler, Phenotypic diversity, population growth, and information in fluctuating environments, Science 309, 2075 (2005).

97 N. Q. Balaban, J. Merrin, R. Chait, L. Kowalik, and S. Leibler, Bacterial persistence as a phenotypic switch, Science 305, 1622 (2004).

98 E. Kussell, R. Kishony, N. Q. Balaban, and S. Leibler, Bacterial persistence: a model of survival in changing environments, Genetics 169, 1807 (2005).

99 P. Visco, R. J. Allen, S. N. Majumdar, and M. R. Evans, Switching and growth for microbial populations in catastrophic responsive environments, Biophys. J. 98, 1099 (2010).

100 P. C. Bressloff, Stochastic switching in biology: from genotype to phenotype, J. Phys. A 50, 133001 (2017).

101 A. Taitelbaum, R. West, M. Assaf, and M. Mobilia, Population dynamics in a changing environment: random versus periodic switching, Phys. Rev. Lett. 125, 048105 (2020).

102 M. L. Ginsberg, Dynamic backtracking, J. Artif. Intell. Res. 1, 25 (1993).

103 M. Luby, A. Sinclair, and D. Zuckerman, Optimal speedup of Las Vegas algorithms, Inf. Process. Lett. 47, 173 (1993).

104 C. P. Gomes, B. Selman, and H. Kautz, Boosting combinatorial search through randomization, Computer Sci. AAAI/IAAI 98, 431 (1998).

105 G. T. Buswell, ”How people look at pictures: a study of the psychology and perception in art”, (Chicago University Press, 1935).

106 D. Noton and L. Stark, Scanpaths in saccadic eye movements while viewing and recognizing patterns, Vision Res. 11, 929 (1971).

107 K. Rayner, Eye movements in reading and information processing, Psychol. Bulletin 85, 618 (1978).

108 K. Rayner, Eye movements in reading and information processing: 20 years of research, Psychol. Bulletin 124, 372 (1998).

109 P. Verghese, Visual search and attention: a signal detection theory approach, Neuron 31, 523 (2001).

110 R. Engbert and R. Kliegl, Microsaccades uncover the orientation of covert attention, Vision Res. 43, 1035 (2003).

111 M. Rolfs, Microsaccades: small steps on a long way, Vision Res. 49, 2415 (2009).

112 R. Engbert, K. Mergenthaler, P. Sinn, and A. Pikovsky, An integrated model of fixational eye movements and microsaccades, Proc. Natl. Acad. Sci. U. S. A. 108, E765 (2011).

113 D. G. Stephen, D. Mirman, J. S. Magnuson, and J. A. Dixon, Lévy-like diffusion in eye movements during spoken-language comprehension, Phys. Rev. E 79, 056114 (2009).

114 C. J. J. Herrmann, R. Metzler, and R. Engbert, A self-avoiding walk with neural delays as a model of fixational eye movements, Sci. Rep. 7, 12958 (2017).

115 P. Blazejczyk and M. Magdziarz, Stochastic modeling of Lévy-like human eye movements, Chaos 31, 043129 (2021).

116 M. P. Eckstein, Visual search: a retrospective, J. Vision 11, 14 (2011).

117 A. Conze and Viswanathan, Path dependent options: the case of lookback options, J. Finance XLVI, 1893 (1991).

118 M. Rubinstein and E. Reiner, Breaking down the barriers, Risk 4, 28 (1991).

119 S. F. Gray and R. E. Whaley, Valuing S&P 500 bear market warrants with a periodic reset, J. Derivatives 5, 99 (1997).

120 W.-Y. Cheng and S. Zhang, The analytics of reset options, J. Derivatives 8, 59 (2000).

121 D. Davydov and V. Linetsky, Pricing and hedging path-dependent options under the CEV process, Manag. Sci. 47, 949 (2001).

122 R. Goerlich, M. Li, S. Albert, G. Manfredi, P.-A. Hervieux, and C. Genet, Noise and ergodic properties of Brownian motion in an optical tweezer: looking at the crossover between Wiener and Ornstein-Uhlenbeck processes, 2007.12246 (2020).

123 S. C. Lim and S. V. Muniandy, Self-similar Gaussian processes for modeling anomalous diffusion, Phys. Rev. E 66, 021114 (2002).

124 F. Thiel and I. M. Sokolov, Scaled Brownian motion as a mean-field model for continuous-time random walks, Phys. Rev. E 89, 012115 (2014).

125 J.-H. Jeon, A. V. Chechkin, and R. Metzler, Scaled Brownian motion: a paradoxical process with a time dependent diffusivity for the description of anomalous diffusion, Phys. Chem. Chem. Phys. 16, 15811 (2014).

126 A. G. Cherstvy and R. Metzler, Ergodicity breaking, ageing, and confinement in generalized diffusion processes with position and time dependent diffusivity, J. Stat. Mech. P05010, (2015).

127 A. Bodrova, A. V. Chechkin, A. G. Cherstvy, and R. Metzler, Ultraslow scaled Brownian motion, New J. Phys. 17, 063038 (2015).

128 H. Safdari, A. G. Cherstvy, A. V. Chechkin, F. Thiel, I. M. Sokolov, and R. Metzler, Quantifying the non-ergodicity of scaled Brownian motion, J. Phys. A 48, 375002 (2015).

129 A. G. Cherstvy, and R. Metzler, Anomalous diffusion in time-fluctuating non-stationary diffusivity landscapes, Phys. Chem. Chem. Phys. 18, 23840 (2016).

130 H. Safdari, A. G. Cherstvy, A. V. Chechkin, A. Bodrova, and R. Metzler, Aging underdamped scaled Brownian motion: Ensemble- and time-averaged particle displacements, nonergodicity, and the failure of the overdamping approximation, Phys. Rev. E 95, 011120 (2017).

131 A. G. Cherstvy, H. Safdari, and R. Metzler, Anomalous diffusion, nonergodicity, and ageing for exponentially and logarithmically time-dependent diffusivity: striking differences for massive versus massless particles, J. Phys. D 54, 195401 (2021).

132 J. L. Lebowitz and O. Penrose, Modern ergodic theory, Phys. Today 26(2), 23 (1973).

133 J.-P. Bouchaud, Weak ergodicity breaking and aging in disordered systems, J. Phys. I 2, 1705 (1992).

134 C. C. Moore, Ergodic theorem, ergodic theory, and statistical mechanics, Proc. Natl. Acad. Sci. U. S. A. 112, 1907 (2015).

135 M. Mangalam and D. G. Kelty-Stephen, Point estimates, Simpsons paradox, and nonergodicity in biological sciences, Neurosci. & Biobehav. Rev. 125, 98 (2021).

136 A. N. Kolmogorov, Wienersche Spiralen und einige andere interessante Kurven im Hilbertschen Raum, C. R. (Doklady) Acad. Sci. URSS (N.S.) 26, 115 (1940).

137 B. B. Mandelbrot and J. W. van Ness, Fractional Brownian motions, fractional noises and applications, SIAM Rev. 10, 422 (1968).

138 F. Biagini, Y. Hu, B. Øksendal, and T. Zhang, ”Stochastic Calculus for Fractional Brownian Motion and Applications”, (Springer-Verlag, London, 2008).

139 W. Deng and E. Barkai, Ergodic properties of fractional Brownian-Langevin motion, Phys. Rev. E 79, 011112 (2009).

140 J.-H. Jeon and R. Metzler, Fractional Brownian motion and motion governed by the fractional Langevin equation in confined geometries, Phys. Rev. E 81, 021103 (2010).

141 J.-H. Jeon and R. Metzler, Inequivalence of time and ensemble averages in ergodic systems: exponential versus power-law relaxation in confinement, Phys. Rev. E 85, 021147 (2012).

142 F. Thiel and I. M. Sokolov, Weak ergodicity breaking in an anomalous diffusion process of mixed origins, Phys. Rev. E 89, 012136 (2014).

143 M. Schwarzl, A. Godec, and R. Metzler, Quantifying nonergodicity of anomalous diffusion with higher order moments, Sci. Rep. 7, 3878 (2017).

144 W. Wang, A. G. Cherstvy, A. V. Chechkin, S. Thapa, F. Seno, X. Liu, and R. Metzler, Fractional Brownian motion with random diffusivity: emerging residual nonergodicity below the correlation time, J. Phys. A 53, 474001 (2020).

145 Y. He, S. Burov, R. Metzler, and E. Barkai, Random time-scale invariant diffusion and transport coefficients, Phys. Rev. Lett. 101, 058101 (2008).

146 A. G. Cherstvy, A. V. Chechkin, and R. Metzler, Anomalous diffusion and ergodicity breaking in heterogeneous diffusion processes, New J. Phys. 15, 083039 (2013).

147 A. G. Cherstvy and R. Metzler, Nonergodicity, fluctuations, and criticality in heterogeneous diffusion processes, Phys. Rev. E 90, 012134 (2014).

148 A. G. Cherstvy, A. V. Chechkin, and R. Metzler, Ageing and confinement in non-ergodic heterogeneous diffusion processes, J. Phys. A 47, 485002 (2014).

149 A. G. Cherstvy and R. Metzler, Ergodicity breaking and particle spreading in noisy heterogeneous diffusion processes, J. Chem. Phys. 142, 144105 (2015).

150 W. Wang, A. G. Cherstvy, X. Liu, and R. Metzler, Anomalous diffusion and nonergodicity for heterogeneous diffusion processes with fractional Gaussian noise, Phys. Rev. E 102, 012146 (2020).

151 H. Hurst, Long term storage capacity of reservoirs, Trans. Am. Soc. Civil Eng. 116, 770 (1951).

152 B. B. Mandelbrot, Harold Edwin Hurst, in “Statisticians of the Centuries”, Eds.: C. C. Heyde et al., (Springer, New York, 2001).

153 J. E. Higham, M. Shahnam, and A. Vaidheeswaran, Anomalous diffusion in a bench-scale pulsed fluidized bed, 2012.05717, (2021).

154 A. W. C. Lau and T. C. Lubensky, State-dependent diffusion: Thermodynamic consistency and its path integral formulation, Phys. Rev. E 76, 011123 (2007).

155 J.-H. Jeon, H. Martinez-Seara Monne, M. Javanainen, and R. Metzler, Anomalous diffusion of phospholipids and cholesterols in a lipid bilayer and its origins, Phys. Rev. Lett. 109, 188103 (2012).

156 J.-H. Jeon, M. Javanainen, H. Martinez-Seara, R. Metzler, and I. Vattulainen, Protein crowding in lipid bilayers gives rise to non-Gaussian anomalous lateral diffusion of phospholipids and proteins, Phys. Rev. X 6, 021006 (2016).

157 R. Metzler, J.-H. Jeon, and A. G. Cherstvy, Non-Brownian diffusion in lipid membranes: experiments and simulations, Biochem. Biophys. Acta BBA-Biomembr. 1858, 2451 (2016).

158 Y. Golan and E. Sherman, Resolving mixed mechanisms of protein subdiffusion at the T cell plasma membrane, Nature Comm. 8, 15851 (2017).

159 T. Sungkaworn, M.-L. Jobin, K. Burnecki, A. Weron, M. J. Lohse, and D. Calebiro, Single-molecule imaging reveals receptor-G protein interactions at cell surface hot spots, Nature 550, 543 (2017).

160 J. Janczura, P. Kowalek, H. Loch-Olszewska, J. Szwabinski, and A. Weron, Classification of particle trajectories in living cells: Machine learning versus statistical testing hypothesis for fractional anomalous diffusion, Phys. Rev. E 102, 032402 (2020).

161 M. Magdziarz, A. Weron, K. Burnecki, and J. Klafter, Fractional Brownian motion versus the continuous-time random walk: a simple test for subdiffusive dynamics, Phys. Rev. Lett. 103, 180602 (2009).

162 J.-H. Jeon, V. Tejedor, S. Burov, E. Barkai, C. Selhuber-Unkel,K. Berg-Sørensen, L. Oddershede, and R. Metzler, In vivo anomalous diffusion and weak ergodicity breaking of lipid granules, Phys. Rev. Lett. 106, 048103 (2011).

163 E. Kepten, I. Bronshtein, and Y. Garini, Ergodicity convergence test suggests telomere motion obeys fractional dynamics, Phys. Rev. E 83, 041919 (2011).

164 K. Burnecki, E. Kepten, J. Janczura, I. Bronshtein, Y. Garini, and A. Weron, Universal algorithm for identification of fractional Brownian motion. A case of telomere subdiffusion, Biophys. J. 103, 1839 (2012).

165 S. C. Weber, M. A. Thompson, W. E. Moerner, A. J. Spakowitz, and J. A. Theriot, Analytical tools to distinguish the effects of localization error, confinement, and medium elasticity on the velocity autocorrelation function, Biophys. J. 102, 2443 (2012).

166 M. P. Backlund, R. Joyner, and W. E. Moerner, Chromosomal locus tracking with proper accounting of static and dynamic errors, Phys. Rev. E 91, 062716 (2015).

167 G. M. Oliveira, A. Oravecz, D. Kobi, M. Maroquenne, K. Bystricky, T. Sexton, and N. Molina, Precise measurements of chromatin diffusion dynamics by modeling using Gaussian processes, bioRxiv: https://doi.org/10.1101/2021.03.16.435699, (2021).

168 A. G. Cherstvy, S. Thapa, C. E. Wagner, and R. Metzler, Non-Gaussian, non-ergodic, and non-Fickian diffusion of tracers in mucin hydrogels, Soft Matter 15, 2481 (2019).

169 I. Y. Wong, M. L. Gardel, D. R. Reichman, E. R. Weeks, M. T. Valentine, A. R. Bausch, and D. A. Weitz, Anomalous diffusion probes microstructure dynamics of entangled F-actin networks, Phys. Rev. Lett. 92, 178101 (2004).

170 M. Levin, G. Bel, and Y. Roichman, Measurements and characterization of the dynamics of tracer particles in an actin network, J. Chem. Phys. 154, 144901 (2021).

171 A. Muralidharan, H. Uitenbroek, and P. E. Boukany, Intracellular transport of electro-transferred DNA cargo is governed by coexisting ergodic and nonergodic anomalous diffusion, submitted (2021). bioRxiv preprint, doi: https://doi.org/10.1101/2021.04.12.435513;

172 A. D. Fernandez, P. Charchar, A. G. Cherstvy, R. Metzler, and M. F. Finnis, The diffusion of doxorubicin drug molecules in silica nanoslits is non-Gaussian, intermittent and anticorrelated, Phys. Chem. Chem. Phys. 22, 27955 (2020).

173 L. Boltzmann, Zur Integration der Diffusionsgleichung bei variabeln Diffusionscoefficienten, Ann. Physik 289, 959 (1894).

174 R. E. Pattle, Diffusion from an instantaneous point source with a concentration-dependent coefficient, Q. J. Mech. Appl. Math. 12, 407 (1959).

175 A. Hansen, E. G. Flekkøy, and B. Baldelli, Anomalous diffusion in systems with concentration-dependent diffusivity: exact solutions and particle simulations, Frontiers Phys. 8, 519624 (2020).

176 A. Hansen, E. G. Flekkøy, and B. Baldelli, Hyperballistic superdiffusion and explosive solutions to the non-Linear diffusion equation, Frontiers Phys. 9, 640560 (2021).

177 A. Fulinski, Communication: How to generate and measure anomalous diffusion in simple systems, J. Chem. Phys. 138, 021101 (2013).

178 M. Heidernätsch, “On the Diffusion in Inhomogeneous Systems”, PhD Thesis (TU Chemnitz, 2015).

179 R. Kazakevicius and J. Ruseckas, Influence of external potentials on heterogeneous diffusion processes, Phys. Rev. E 94, 032109 (2016).

180 X. Wang, W. Deng, and Y. Chen, Ergodic properties of heterogeneous diffusion processes in a potential well, J. Chem. Phys. 150, 164121 (2019).

181 M. A. F. dos Santos, V. Dornelas, E. H. Colombo, and C. Anteneodo, Critical patch size reduction by heterogeneous diffusion, Phys. Rev. E 102, 042139 (2020).

182 T. Sandev, A. Schulz, H. Kantz, and A. Iomin, Heterogeneous diffusion in comb and fractal grid structures, Chaos, Solitons & Fractals 114, 551 (2018).

183 N. Leibovich and E. Barkai, Infinite ergodic theory for heterogeneous diffusion processes, Phys. Rev. E 99, 042138 (2019).

184 R. Kazakevicius and A. Kononovicius, Anomalous diffusion in nonlinear transformations of the noisy voter model, Phys. Rev. E 103, 032154 (2021).

185 S. K. Ghosh, A. G. Cherstvy, D. S. Grebenkov, and R. Metzler, Anomalous, non-Gaussian tracer diffusion in crowded two-dimensional environments, New J. Phys. 18, 013027 (2016).

186 P. Massignan, C. Manzo, J. A. Torreno-Pina, M. F. Garcia-Parajo, M. Lewenstein, and G. J. Lapeyre Jr.,, Nonergodic subdiffusion from Brownian motion in an inhomogeneous medium, Phys. Rev. Lett. 112, 150603 (2014).

187 R. Jain and K. L. Sebastian, Diffusion in a crowded, rearranging environment, J. Phys. Chem. B 120, 3988 (2016).

188 H. Berry and H. Chate, Anomalous diffusion due to hindering by mobile obstacles undergoing Brownian motion or Ornstein-Uhlenbeck processes, Phys. Rev. E 89, 022708 (2014).

189 F. Trovato and V. Tozzini, Diffusion within the cytoplasm: a mesoscale model of interacting macromolecules, Biophys. J. 107, 2579 (2014).

190 M. Weiss, Crowding, diffusion, and biochemical reactions, Intl. Rev. Cell & Mol. Biol. 307, 383 (2014).

191 M. J. Saxton, Wanted: scalable tracers for diffusion measurements, J. Phys. Chem. B 118, 12805 (2014).

192 A. Russian, M. Dentz, and P. Gouze, Self-averaging and weak ergodicity breaking of diffusion in heterogeneous media, Phys. Rev. E 96, 022156 (2017).

193 Y. Lanoiselee, N. Moutal, D. S. Grebenkov, Diffusion-limited reactions in dynamic heterogeneous media, Nature Comm. 9, 4398 (2018).

194 S. Basak, S. Sengupta, and K. Chattopadhyay, Understanding biochemical processes in the presence of sub-diffusive behavior of biomolecules in solution and living cells, Biophys. Rev. 11, 851 (2019).

195 P. Witzel, M. Götz, Y. Lanoiselee, T. Franosch, D. S. Grebenkov, and D. Heinrich, Heterogeneities shape passive intracellular transport, Biophys. J. 117, 203 (2019).

196 G. Munoz-Gil, M. A. Garcia-March, C. Manzo, A. Celi, and M. Lewenstein, Diffusion through a network of compartments separated by partially-transmitting boundaries, Front. Phys. 7, 31 (2019).

197 A. Sabri, X. Xu, D. Krapf, and M. Weiss, Elucidating the origin of heterogeneous anomalous diffusion in the cytoplasm of mammalian cells, Phys. Rev. Lett. 125, 058101 (2020).

198 I. Chakraborty and Y. Roichman, Disorder-induced Fickian, yet non-Gaussian diffusion in heterogeneous media, Phys. Rev. Res. 2, 022020(R) (2020).

199 E. B. Postnikov, A. V. Chechkin, and I. M. Sokolov, Brownian yet non-Gaussian diffusion in heterogeneous media: from superstatistics to homogenization, New J. Phys. 22, 063046 (2020).

200 M. J. Skaug and D. K. Schwartz, Tracking nanoparticle diffusion in porous filtration media, Ind. Eng. Chem. Res. 54, 4414 (2015).

201 M. J. Skaug, L. Wang, Y. Ding, and D. K. Schwartz, Hindered nanoparticle diffusion and void accessibility in a three-dimensional porous medium, ACS Nano 9, 2148 (2015).

202 R. Sarfati, C. P. Calderon, and D. K. Schwartz, Enhanced diffusive transport in fluctuating porous media, ACS Nano 15, 7392 (2021).

203 C. B. Mast, S. Schink, U. Gerland, and D. Braun, Escalation of polymerization in a thermal gradient, Proc. Natl. Acad. Sci. 110, 8030 (2013).

204 D. Breoni, H. Löwen, and R. Blossey, Active noise-driven particles under space-dependent friction in one dimension, 2102.09944 (2021).

205 P. Massignan, A. Lampo, J. Wehr, and M. Lewenstein, Quantum Brownian motion with inhomogeneous damping and diffusion, Phys. Rev. A 91, 033627 (2015)

206 A. Geraldi, S. De, A. Laneve, S. Barkhofen, J. Sperling, P. Mataloni, and C. Silberhorn, Transient subdiffusion via disordered quantum walks, Phys. Rev. Res. 3, 023052 (2021).

207 J. Crank, ”The Mathematics of Diffusion”, (Oxford University Press, 1975).

208 W. S. Gurney and R. M. Nisbet, The regulation of inhomogeneous populations, J. Theor. Biol. 52, 441 (1975).

209 M. E. Gurtin and R. C. MacCamy, On the diffusion of biological populations. Math. Biosci. 33, 35 (1977).

210 W. I. Newman, Some exact solutions to a non-linear diffusion problem in population genetics and combustion, J. Theor. Biol. 85, 325 (1980).

211 D. G. Aronson, “The porous medium equation”, in “Nonlinear Diffusion Problems”, pp. 1–46, in “Lecture Notes in Mathematics”, vol. 1224, Eds.: A. Dold and B. Eckmann (Springer-Verlag, 1986).

212 I. C. Christov and H. A. Stone, Resolving a paradox of anomalous scalings in the diffusion of granular materials, Proc. Natl. Acad. Sci. U. S. A. 109, 16012 (2012).

213 metaphorically, after Henri de Pitot (also known as “satellite droplet(s)”), see, e.g., Ya. E. Gegusin, ”The Drop”, (Moscow, “Nauka”, 1973). [in Russian]

214 A. G. Cherstvy, S. Thapa, Y. Mardoukhi, A. V. Chechkin, and R. Metzler, Time averages and their statistical variation for the Ornstein-Uhlenbeck process: role of initial particle conditions and relaxation to stationarity, Phys. Rev. E 98, 022134 (2018).

215 B. Goswami, N. Boers, A. Rheinwalt, N. Marwan, J. Heitzig, S. F. M. Breitenbach, and J. Kurths, Abrupt transitions in time series with uncertainties, Nature Comm. 9, 48 (2018).

216 R. Hou, A. G. Cherstvy, R. Metzler, and T. Akimoto, Biased continuous-time random walks for ordinary and equilibrium cases: facilitation of diffusion, ergodicity breaking and ageing, Phys. Chem. Chem. Phys. 20, 20827 (2018).

217 V. Zaburdaev, S. Denisov, and J. Klafter, Lévy walks, Rev. Mod. Phys. 87, 483 (2015).

218 G. E. Uhlenbeck and L. S. Ornstein, On the theory of the Brownian motion, Phys. Rev. 36, 823 (1930).

219 A. G. Cherstvy, D. Vinod, E. Aghion, A. V. Chechkin, and R. Metzler, Time averaging, ageing and delay analysis of financial time series, New J. Phys. 19, 063045 (2017).

220 S. Ritschel, A. G. Cherstvy, and R. Metzler, Universality of delay-time averages for financial time series: analytical results, computer simulations, and analysis of historical stock-market prices, submitted, (2021).

221 D. Vinod, A. G. Cherstvy, I. M. Sokolov, and R. Metzler, Resetting, time-averaging, and nonergodicity for geometric Brownian motion, work in preparation, (2021).

222 A. G. Cherstvy, D. Vinod, E. Aghion, I. M. Sokolov, and R. Metzler, Scaled geometric Brownian motion features sub-or superexponential ensemble-but linear time-averaged mean-squared displacements, submitted, (2021).

223 V. Stojkoski, T. Sandev, L. Kocarev, and A. Pal, Geometric Brownian motion under stochastic resetting: a stationary yet non-ergodic process, 2104.01571 (2021).

224 S. Petrovskii and A. Morozov, Dispersal in a statistically structured population: fat tails revisited, Am. Naturalist 173, 278 (2009).

225 S. Hapca, J. W. Crawford, and L. M. Young, Anomalous diffusion of heterogeneous populations characterized by normal diffusion at the individual level, J. Royal Soc. Interface 6, 111 (2009).

226 M. V. Chubynsky and G. W. Slater, Diffusing diffusivity: a model for anomalous, yet Brownian, diffusion, Phys. Rev. Lett. 113, 098302 (2014).

227 T. Uneyama, T. Miyaguchi, and T. Akimoto, Fluctuation analysis of time-averaged mean-square displacement for the Langevin equation with time-dependent and fluctuating diffusivity, Phys. Rev. E 92, 032140 (2015).

228 A. V. Chechkin, F. Seno, R. Metzler, and I. M. Sokolov, Brownian yet non-Gaussian diffusion: from superstatistics to subordination of diffusing diffusivities, Phys. Rev. X 7, 021002 (2017).

229 R. Jain and K. L. Sebastian, Lévy flight with absorption: a model for diffusing diffusivity with long tails, Phys. Rev. E 95, 032135 (2017).

230 A. Weron, K. Burnecki, E. J. Akin, L. Sole, M. Balcerek, M. M. Tamkun, and D. Krapf, Ergodicity breaking on the neuronal surface emerges from random switching between diffusive states, Sci. Rep. 7, 5404 (2017).

231 T. Miyaguchi, T. Uneyama, and T. Akimoto, Brownian motion with alternately fluctuating diffusivity: stretched-exponential and power-law relaxation, Phys. Rev. E 100, 012116 (2019).

